# Updating the RZooRoH package for the analysis of inbreeding, identity-by-descent and relatedness from genomic data

**DOI:** 10.1101/2025.07.08.663647

**Authors:** Natalia S Forneris, Pierre Faux, Mathieu Gautier, Tom Druet

**Affiliations:** Unit of Animal Genomics, GIGA-R & Faculty of Veterinary Medicine, University of Liège, Liège, Belgium; GenPhySE, Université de Toulouse, INRAE, ENVT, 31326 Castanet Tolosan, France; CBGP, INRAE, CIRAD, IRD, L’institut Agro, Université de Montpellier, Montpellier, France

**Keywords:** Inbreeding, homozygosity-by-descent (HBD), ROH, identity-by-descent (IBD), kinship

## Abstract

The RZooRoH R package was implemented to characterize individual inbreeding levels. It identifies DNA segments inherited twice from a common ancestor through different paths, which are known as homozygous-by-descent (HBD) segments. The package accepts different data formats and provides multiple outputs: HBD segments, inbreeding rates and genome-wide and locus-specific HBD probabilities. In addition, it partitions HBD levels into multiple HBD classes. The length distribution varies between these classes, which therefore correspond to distinct groups of ancestors that can be traced back to different generations in the past. This provides information about mating structure and recent demographic history. The computational performance of the package has been substantially improved, enabling, for example, computing times to be reduced when working with whole-genome sequence data and more HBD classes to be fitted. It is now possible to fit one class per past generation, which facilitates interpretation of the results. Since we have previously demonstrated that the ZooRoH model can be used to characterize identity-by-descent (IBD) between haploid individuals or phased haplotypes, this option has been included in the new package version. Estimating kinship by characterizing IBD levels between the four possible pairs of haplotypes from two individuals is another feature we added to the package. Finally, new options allow models to be refined, for instance by defining HBD classes as intervals or constant inbreeding rates for neighboring classes. Overall, the new version of the package offers improved computational efficiency and interpretability when characterizing inbreeding, IBD and relatedness levels.

## 1 INTRODUCTION

The RZooRoH package (Bertrand et al., 2019) implements a model-based approach developed to characterize individual inbreeding levels from genomic data by estimating homozygous-by-descent (HBD) probabilities and identifying HBD segments along their genomes (Druet & Gautier, 2017, 2022). HBD segments occur when an individual inherits the same chromosomal segment twice from an ancestor, and are more common in inbred populations. They typically appear as long stretches of homozygous markers called runs-of-homozygosity (RoH). HBD probabilities and HBD segments are useful in many applications. Indeed, the proportion of the genome in HBD segments is an estimator of the inbreeding coefficient (e.g., Leutenegger et al., 2003; McQuillan et al., 2008; Druet & Gautier, 2017), a measure that can be used to manage populations with small effective population size or to study inbreeding depression. In addition, as they provide locus-specific information, they allow so-called homozygosity mapping experiments to identify recessive variants associated with disease (Seelow et al., 2009) or to identify genetic variants associated with inbreeding depression. Their length is indicative of the number of generations to the common ancestor, as segments associated with more distant ancestors are expected to be shorter because recombination has more opportunities to cut the transmitted segment. Therefore, the length-based classification of HBD segments is informative about the recent demographic history of a population (Kirin et al., 2010) and can be used to study whether recent inbreeding is more deleterious (Szpiech et al., 2013; Stoffel et al., 2021; Naji et al., 2024).

An important feature of RZooRoH is that it relies on a model-based approach that uses allele frequencies, genotyping error rates and genetic distances to estimate the proportion of the genome associated with multiple HBD classes. Each class is defined by its rate parameter, which determines the expected length of HBD segments in the class. Different classes therefore correspond to different groups of ancestors present in different past generations. For example, low-rate HBD classes capture the longer HBD segments associated with more recent common ancestors. The model-based approach allows the use of different types of genetic data, including genotypes, genotype probabilities and likelihoods, and allele counts. The approach has already been evaluated and compared with other approaches in several studies (e.g., Druet & Gautier, 2017, 2022; Solé et al., 2017; Alemu et al., 2021; Duntsch et al., 2021; Lavanchy & Goudet, 2023; Forneris et al., 2025). The model-based approach is particularly valuable when data are heterogeneous (e.g. variable genotyping error rate, variable marker spacing) or with reduced information (low-marker density and/or sequencing coverage). This is typically the case when analysing genomic data generated with exome-sequencing (Magi et al., 2014), genotyping-by-sequencing, low-fold sequencing or from low density marker arrays. Consequently, the package is used in livestock populations (e.g., Solé et al., 2017; Bhati et al., 2020; Moyse et al., 2022; Zhang et al., 2022) and increasingly in conservation genetics (e.g., Druet et al., 2020; Lobo et al., 2023), or for the study of feral (e.g., Gautier et al., 2024; Thompson et al., 2024) and wild populations (Coimbra et al., 2021; Pacheco et al., 2022; Nigenda-Morales et al., 2023).

We herein present significant updates to the RZooRoH package (v0.4.0) including the implementation of a new algorithm that can dramatically reduce computational times and the definition of new HBD classes defined by a range of rates of coancestry changes, facilitating interpretation of results. In addition, as it has been previously shown that the model can also be used to estimate identity-by-descent (IBD) between pairs of haplotypes (Forneris et al., 2025), we have also extended the model to IBD and relatedness analysis. Importantly, an updated vignette provides a full description of all the new implementations.

## 2 THE RZOOROH PACKAGE AND ITS NEW FEATURES

The RZooRoH package was originally implemented to identify HBD segments and characterize inbreeding levels using a model-based approach (Bertrand et al., 2019). At the heart of the ZooRoH model is a hidden Markov model (HMM) that describes individual genomes as mosaics of HBD and non-HBD segments - the two hidden states of the model. The key components of this HMM are the initial state probabilities, transition probabilities, and emission probabilities. The initial state probabilities represent the probability of beginning a sequence in a specific state and also correspond to the stationary distribution. The transition probabilities define the probability of moving from one state to another between successive marker positions. They depend on the probability of a segment ending, as well as on the frequency of HBD segments, denoted *ρ*. The probability of a segment ending is obtained from the exponential distribution as 1 − 𝑒^Rd^, where *d* is the genetic distance between the markers (in Morgans) and R is the rate parameter from the exponential distribution. This rate is also referred to as the rate of coancestry change, since it determines the rate at which we transition from HBD to non-HBD (and vice versa), which implies a change in ancestry. In this model, the length of HBD segments follows an exponential distribution with a rate of R, giving a mean of 1/R Morgans or 100/R cM. Finally, the emission probabilities correspond to the probability of observing the genotype or sequence data given the underlying state. In HBD segments, homozygosity is expected, whereas in non-HBD segments, genotypes follow Hardy-Weinberg proportions. These emission probabilities account for allele frequencies, genotyping error rates, and genotype uncertainty when working with low-fold sequence data, for example. The simplest model has two states, similar to models described by Leutenegger et al. (2003), Narasimhan et al. (2016) and Vieira et al. (2016), and assumes that all HBD segments have the same expected length. In this case, the parameter *ρ*, which defines the frequency of HBD segments, is also an estimator of the inbreeding coefficient *F*. A key feature of the ZooRoH model is the definition of multiple HBD classes. Each class has its own rate of coancestry change, denoted R_c_ (where *c* is the index of the class), and the expected length of HBD segments varies accordingly. More precisely, the length distribution remains exponential, but with different means equal to 100/R_c_ cM. This enables the contribution of different groups of ancestors to be partitioned into different HBD classes. These HBD classes are defined as successive layers of ancestors as described in Figure 1 (Druet & Gautier, 2022). Within each layer, the genome is modelled as a mosaic of HBD and non-HBD segments with a single rate of coancestry change. The non-HBD class is then modelled as a mosaic of HBD and non-HBD segments, with higher rates of coancestry change corresponding to more distant ancestors (i.e., having more remote ancestors means there are more frequent ancestry changes, resulting in shorter HBD and non-HBD segments). In such a model, the frequency of HBD segments varies between layers, with each HBD class having its own parameter ρ_c_ (where *c* is the class index). This parameter corresponds to the inbreeding rate in the layer (see Druet & Gautier, 2022), i.e. the relative proportion of new inbreeding accumulated in the layer. Accordingly, the ρ_c_ parameters will be referred to as the ‘inbreeding rates per layer’. To facilitate understanding of the model, Figure 1 describes its main features and the properties of HBD segments.

**Figure 1.**
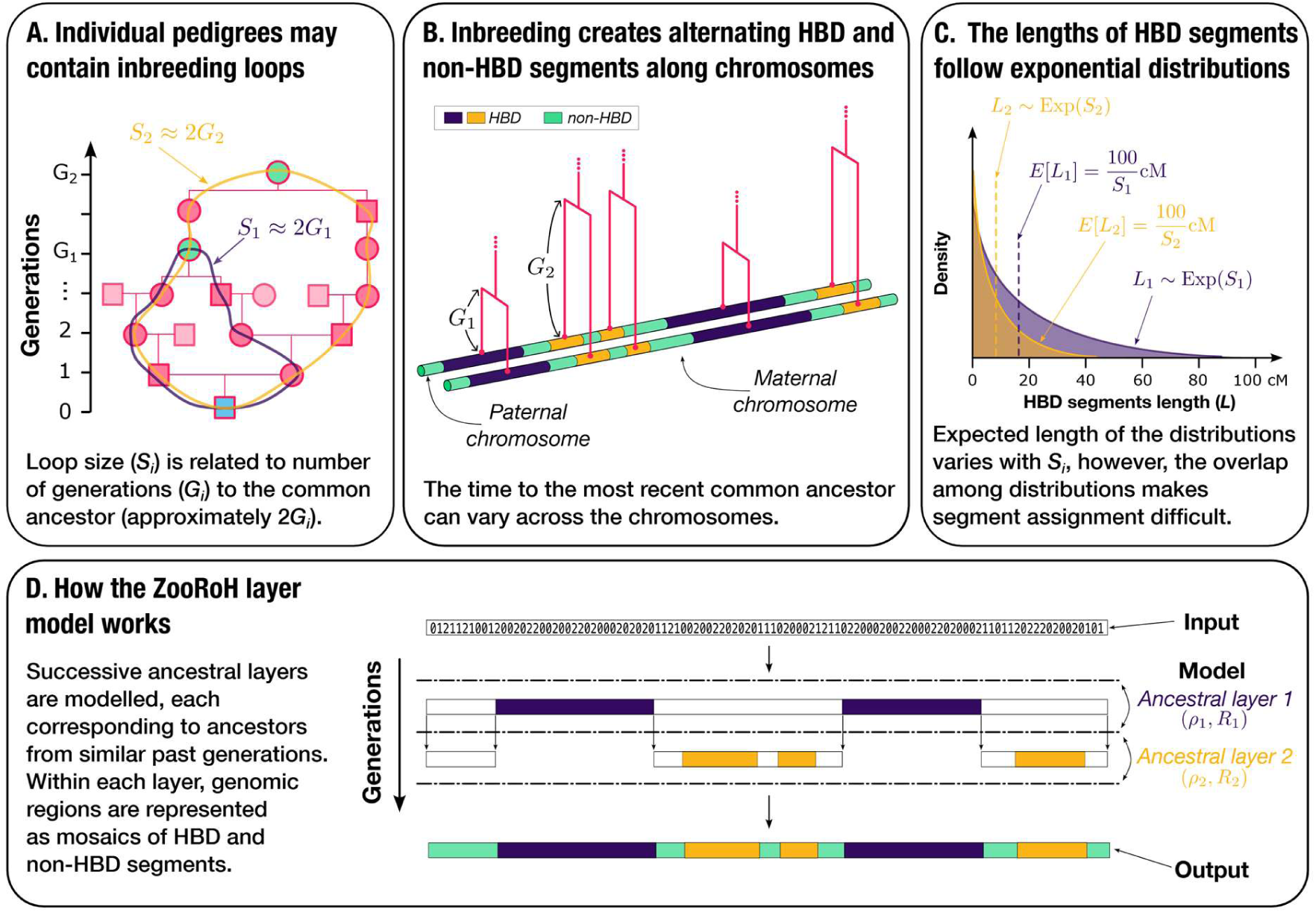
Description of the homozygous-by-descent (HBD) process and the ZooRoH model. (**A**) Inbreeding occurs when an individual’s parents are related, i.e. when they are connected in a pedigree (as represented by the two loops connecting the parents of the focal individual). Inbreeding levels are a function of the number and size S_i_ of the inbreeding loops (i.e. the number of meiosis or parent-offspring connections in the loop). In idealized cases, the size of an inbreeding loop is equal to twice the number of generations to the common ancestor (G_i_). (**B**) Inbreeding generates HBD segments along chromosomes, which are regions of DNA that are inherited from the same common ancestor. These segments alternate with non-HBD segments. HBD segments may trace back to different ancestors, who may belong to different past generations. (**C**) The length of HBD segments in Morgans is exponentially distributed with a rate parameter equal to S_i_, the size of the underlying inbreeding loop. Therefore, the expected length is 1/S_i_ Morgans (or 100/S_i_ cM), although the distribution exhibits high variability. (**D**) The ZooRoH model consists of a hidden Markov model with multiple HBD classes. It models individual genomes as a mosaic of HBD and non-HBD segments, taking as input genotypes or genotype likelihoods. The model considers successive nested ancestral layers (groups of ancestors present in approximately the same past generations). Within each layer, genomic regions are modelled as a mosaic of HBD and non-HBD segments. These are then modelled as a mosaic of HBD and non-HBD segments in the next layer, which corresponds to more distant ancestors. The layer-specific parameter ρ_c_ specifies the frequency of HBD segments within layer *c* and corresponds to the inbreeding rate per layer (i.e. the relative increase in inbreeding within the layer). The length of HBD segments originating from layer *c* is modelled using an exponential distribution with rate R_c_, called the rate of coancestry change within layer *c* (i.e. the frequency at which segments end and new ones start). The length distribution in layer *c* (or HBD class) matches that associated with inbreeding loops of size R_c_. Note that each layer is defined by a single rate of coancestry change, which is common to all ancestors captured by the layer. Hence, the boundaries between layers (in terms of generation captured) are not clearly defined. The output comprises the partitioning of the genome into different HBD classes (i.e. the mosaic of HBD and non-HBD segments), the probability of each position in the genome belonging to the different HBD (and non-HBD) classes, and the identified HBD segments.

The RZooRoH package implements three main functions: zoodata(), zoomodel() and zoorun(). The first function reads data in several formats, including genotypes, genotype probabilities, genotype likelihoods and read counts. The second function is used to define the model, including the number of HBD classes, their rate of coancestry change R_c_ and the initial values of the inbreeding rates per layer ρ_c_. The last function runs the defined model on the data and can be applied in parallel. The main outputs are the estimated parameters, including the parameters ρ_c_, and R_c_ (if not fixed). Other outputs include the estimated genome-wide and locus-specific HBD probabilities per class (provided by the forward–backward algorithm) and the identified HBD segments (obtained using the Viterbi algorithm). Additionally, the package provides several plotting and accessor functions to facilitate data manipulation. The following paragraphs outline key updates to the package, including a new function for estimating allele frequencies and additional formats for the zoodata() function, new modelling options for the zoomodel() function, an enhanced algorithm for the zoorun() function and a new zookin() function for estimating relatedness.

### 2.1 A new decoding algorithm for highly improved runtime performance

The layer model described above shares several features with the Pairwise Sequentially Markov Coalescent (PSMC) model (Li & Durbin, 2011) or SMC models when applied to two haplotypes (Sheehan et al., 2013). Harris et al. (2014) proposed a faster decoding algorithm for such models that depends linearly on the number of HBD classes, rather than quadratically. To reduce RZooRoH runtime, this algorithm has been implemented in the forward algorithm to estimate the likelihood of the model for a set of parameters (used for parameter estimation), and in the forward-backward algorithm to estimate local and global HBD probabilities. This important new feature enables RZooRoH to scale to larger cohorts and process sequenced individuals faster, thereby reducing the overall computational resource usage. Runtime reduction is particularly important when estimating relatedness (see below), as the model is applied to all pairs of individuals four times. It also allows many more HBD classes to be fitted in reasonable time, offering a finer partitioning of autozygosity and enabling more subtle differences between groups of individuals to be observed. At the most extreme, this gives also the possibility to fit one HBD class per past generation of ancestors, improving interpretability (see below).

### 2.2 Extension to the analysis of haploid pairs

The ZooRoH model was originally developed to model HBD between two haplotypes within the same individual. However, the same model can be applied to any pair of haplotypes as long as they are known. This is noticeably the case for haploid individuals or for gonosomes in the heterogametic sex (e.g., males X or female Z chromosomes). In fact, when a single IBD class is fitted, ZooRoH is very similar to an HMM that has been specifically developed for haploid organisms, called hmmIBD (Schaffner et al., 2018). In diploid individuals, this approach requires phasing, which is mostly done with statistical approaches and can introduce errors. Nevertheless, Forneris et al. (2025) showed that using ZooRoH on phased haplotypes was efficient for estimating both genome-wide and locus-specific IBD levels. Accordingly, we added the ability to read haploid data with the zoodata() function and to perform IBD analysis by setting ‘ibd’ to TRUE with the zoorun() function. We have also added two data formats: phased VCF files and the so-called ‘haps’ format. With this new option, the outputs are equivalent to those obtained with HBD analyses, except that the results are obtained for pairs of haplotypes, and HBD is replaced by IBD (i.e. the probability of sharing IBD segments, the identification of IBD segments and their partitioning into different classes defined by their rates of coancestry change R_c_).

Overall, this new option allows to apply the multiple HBD classes model to haploid organisms. This allows IBD tracts between pairs of haplotypes to be partitioned into different length-based classes (separating recent from ancient relatedness), offering additional information on demographic history and relatedness, for example. Furthermore, this new function is required for estimating relatedness using the ZooRoH model. However, it is not intended to replace other tools used for identifying IBD segments in diploid organisms.

### 2.3 Estimation of relatedness

IBD analysis among the four possible pairs of haplotypes from two individuals (each pair consisting of one haplotype from each individual) allows estimation of their relatedness. Forneris et al. (2025) showed that using this approach with the ZooRoH model provided an accurate estimator of relatedness. In addition, the relatedness is then partitioned into multiple IBD classes and is informative about how distantly related two individuals are (in terms of generations). Therefore, this approach has been implemented as a new function called zookin(), to facilitate its application by the users. With this analysis, the outputs are estimators of kinship levels, partitioned into different age-related IBD classes, and identification of IBD segments shared among the four parental haplotypes. Additionally, the probability that two randomly selected haplotypes, one from each individual, are IBD is provided for each marker position. These probabilities reflect the local kinship between these individuals and correspond to the expected level of inbreeding in their (future) offspring (see Forneris et al., 2025).

Using RZooRoH to estimate relatedness is useful for breeding management, as Forneris et al. (2025) showed that it can accurately predict the expected level of inbreeding for different matings. They also demonstrated that it is an accurate estimator of relatedness for relatively recent base populations (e.g. 15 or 50 generations in the past) when compared with other methods. Therefore, this measure of relatedness could be used to manage diversity in populations with reduced effective population size (N_e_). The classification into different IBD categories should also facilitate interpretation of relationships and distinguish between recent and background relatedness. Finally, relatedness estimated with RZooRoH can be directly compared with HBD levels estimated using the same model. Such comparisons between inbreeding and kinship can be used to test whether matings are random, whether there is population structure, or whether inbred individuals are less likely to mate together (inbreeding avoidance). Overall, the RZooRoH relatedness measure is most useful in populations with lower N_e_ and presenting some inbreeding.

### 2.4 New specific modelling features

#### 2.4.1 Constant inbreeding rate for groups of successive layers (zoomodel ‘Step’ option)

In certain situations, users may choose to define models with a large number of HBD classes. For instance, fitting a model with 50 layers - with their rates of coancestry changes corresponding to all even numbers between 2 and 100 - would represent the contribution of the last 50 generations of ancestors to HBD in an idealized population, with one layer representing each generation. Indeed, ancestors present G generations in the past would generate HBD segments of an exponentially distributed length with a rate of 2G (see Figure 1). Such a model would enable a more precise identification of the generations contributing to HBD and allow inbreeding rates to be estimated on a per-generation basis, making the results easier to interpret. For example, this approach would be useful when all inbreeding is associated with a brief, intense demographic event (e.g. a bottleneck), as it provides information about the timing of this event. However, this would result in a large number of parameters needing to be optimized, which would increase computing times. Some optimization algorithms rely indeed on estimating the likelihood after making small sequential changes to each parameter, which requires much more exploration and iterations when the number of parameters is high. By forcing groups of successive layers to have identical inbreeding rates, the number of parameters is reduced to one rate per group, thereby reducing the number of iterations needed for convergence. This also reduces the complexity of the problem, leading to fewer computational issues. Furthermore, this relationship between inbreeding rates from neighboring layers may better fit the data, as inbreeding rates are expected to display some autocorrelation, i.e. without abrupt changes between successive layers. Therefore, we implemented a new option in the zoorun() function called ‘Step’, which allows the inbreeding rates per layer to be fitted with a step function (i.e. modelling constant inbreeding rates for groups of successive layers), while retaining the advantages of a model with more HBD classes, as described above.

#### 2.4.2 Refinement of HBD classes boundaries (zoomodel ‘Interval’ option)

The rate of coancestry changes are associated to the number of generations to the common ancestor of an HBD segment. In the default ZooRoH model, a single rate of coancestry change is defined per HBD class or ancestral layer (see Figure 1). This parameter corresponds to the global rate of coancestry change for all generations of ancestors captured by that HBD class. However, boundaries between the HBD classes are not clearly defined and it remains difficult to determine exactly which generations of ancestors are captured by each HBD class. Knowing the HBD class boundaries would provide more explicit information about the past generations captured by each HBD class, as well as the number of generations captured, which could help estimate the inbreeding rate per generation. Assuming that ancestors occur in non-overlapping generations, indexed by integers from 1 to G_N_ (where G_N_ is the most ancient generation captured), and that the lengths of inbreeding loops are equal to twice the number of generations to the ancestors, the rate of coancestry changes take on even values ranging from 2 to 2G_N_. In this framework, each HBD class can be represented as a set of generations of ancestors bounded between G_c_ and G_c+1_ (where c is the index of the HBD class), with rates of coancestry change ranging from 2G_c_ to 2G_c+1_. In this view, each of these multigenerational HBD classes contains all single-generation HBD classes within the selected boundaries. While such a model would facilitate interpretation (see above), it would come with higher computational complexities. For example, transition probabilities would need to account for transitions between any generation within a class and any other generation, and the probability of belonging to a given HBD class would be obtained by summing the probabilities across all generations included in that class. We relied on the computations described in the appendix from Harris et al. (2014) to implement such an approach. It involves some approximations compared to a model that fits all generations separately, but has the advantage of defining fewer classes and thus remains more tractable, particularly if the user wishes to include more ancient generations. This new option can be used when defining the model with the zoomodel() function and by setting ‘HBDclass’ to ‘Interval’.

Nevertheless, this modelling remains computationally demanding. As some computations are performed for each marker interval and depend on the genetic distance between markers, we have added an option to perform them for a finite set of about 500 possible distances, chosen so that the difference between the true and the rounded recombination rate is always within 10% of the true value. This approach speeds up the computations and becomes more advantageous as the number of markers increases. This ‘RecTable’ option can be selected when calling the zoorun() function.

### 2.5 Additional updates

The ZooRoH model takes allele frequencies into account. Earlier versions estimated these frequencies using simple methods (alternatively, the user could provide them). While these approaches were efficient, more precise estimates could be obtained when genotypes are not unambiguously called (e.g. when genotype probabilities have intermediate values). Therefore, to achieve this, we have also added the possibility to estimate allele frequencies using the EM algorithm described by Kim et al. (2011) that takes this uncertainty into account.

In some relatively rare cases, we observed that the optimization function used to estimate parameters either returned errors or failed to converge. Therefore, when calling zoorun(), we added options for users to select alternative optimisation functions, change the number of iterations or impose constraints on the parameters, in the hope that these different settings would improve convergence.

Finally, new accessor functions were implemented to make it easier to extract and plot the results, and to merge or update results obtained in different runs (‘merge_zres’ and ‘update_zres’).

## 3 MATERIAL AND METHODS

### 3.1 Data

#### 3.1.1 Real Datasets

To illustrate the new features of the package, we reanalyzed real datasets from the Amsterdam Island feral cattle (Gautier et al., 2024), European bison (Druet et al., 2020) and honey bees (Leroy et al., 2024). For the cattle population (referred to as TAF after Gautier et al. (2024)), we had 18 individuals genotyped on a medium-density (MD) array (23,679 polymorphic SNPs), of which eight were also whole-genome sequenced (WGS) (9,036,293 polymorphic SNPs after filtering). We also extracted from the sequence data 666,962 polymorphic SNPs present on commercial bovine genotyping arrays, corresponding to a high density (HD) array. The dataset and SNP filtering are fully described in Gautier et al. (2024). For European bison, we selected individuals from the lowland line (denoted WL following Druet et al. (2020)) genotyped with the BovineHD array. After SNP filtering, these individuals had genotypes at 15,673 autosomal polymorphic SNPs, corresponding to a low-density genotyping array. Finally, we selected the black honey bee population from the Ouessant island to illustrate applications on haploid data. Original WGS data from Wragg et al. (2022) was available for 40 haploid drones. We extracted these individuals from the dataset used by Leroy et al. (2024) and kept bi-allelic SNPs with a QUAL > 1000, without heterozygous calls and with a call rate above 95% in the population, resulting in 772,495 polymorphic SNPs.

#### 3.1.2 Simulated datasets

We used datasets that were simulated using SLiM 4.0.1. (Haller & Messer, 2023) and msprime 1.2.0 (Baumdicker et al., 2022), in order to evaluate the ability of the new relatedness estimator to distinguish between recent and background relatedness. We simulated a population that experienced an extreme bottleneck (N_e_ = 10) 15–20 generations ago, followed by recovery to a population size of over 1,000 individuals. This scenario was designed to mimic the feral cattle population. Secondly, we simulated a population with a relatively small effective population size (N_e_ = 250) over the last 1,000 generations. This results in lower background relatedness levels spread over more generations. In both cases, the genome consisted of 25 chromosome pairs, each 100 cM long, and a subset of 25,000 SNPs was selected. The details of these simulations are provided in Table S1, including N_e_ evolution across generations and the final number of individuals. Each scenario was repeated five times.

### 3.2 Applications and evaluation

#### 3.2.1 Computing time

To quantify the speed increase obtained with the newly implemented algorithm, we compared the runtime of versions 0.4.0 and 0.3.2.1 of the package on the TAF datasets at different marker densities. At the WGS, HD and MD levels, we fitted models with 15, 13 and 10 HBD classes, respectively, with the default rates of coancestry change (R_c_ = 2^c^, where c is the class index). Additionally, at the MD level, we applied a model with 50 HBD classes, with R_c_ taking all even values between 2 and 100. These computing times were evaluated on a computing cluster equipped with compute nodes with two 32 cores AMD Epyc Rome 7542 CPUs running at 2.9 GHz. We also reported the memory usage.

#### 3.2.2 IBD analysis and partitioning in pairs of haplotypes: application to a haploid organism

To illustrate the application of the haploid function to a haploid organism, we used it with black honey bee data. We estimated pairwise IBD among the 40 haploid bee drones (780 pairs). First, we ran a model with a single HBD class, estimating the rate of coancestry change, to demonstrate the equivalence of this model to the model implemented in hmmIBD (Schaffner et al., 2018), which is a reference method for haploid organisms. We then ran a 14-layer model with the default rates of coancestry change (R_c_ = 2^c^, where c is the class index) to demonstrate the advantages of a multiple HBD class model over a single HBD class model. For these analyses, we assumed a constant recombination rate of 22.67 cM/Mb (e.g., Webster, 2019).

#### 3.2.3 Relatedness estimation

To illustrate the properties of the new relatedness estimator, we used a nine-layer model with default rates of coancestry changes to estimate and partition relatedness among the 18 genotyped TAF individuals. We show how the output can be used to partition kinship in different IBD classes, thereby facilitating interpretation. We also show that function provides IBD estimation along the chromosomes, at that the partitioning approach can be applied to these IBD values too.

To further evaluate the function’s ability to distinguish between recent and background relatedness, we applied it to the simulated datasets using the same model. First, we phased the entire sample of each repetition with Beagle 5.5 (Browning et al., 2021). Then, we selected pairs of individuals with specific relationships, such as parent–offspring, half-siblings, half uncle-nephew and half-cousins, as well as unrelated individuals (no common ancestor in the last three generations). These pair types were selected to obtain a sufficient number of pairs, and because they present both recent and background relatedness (50 pairs per relationship and replicate were selected). The expected levels of recent relatedness. The results were compared with those obtained using KING (Manichaikul et al., 2010), a popular kinship estimation tool that can correct for background relatedness and also proposes an estimator based on IBD segments. To assess the impact of phasing errors, we repeated the analyses, this time phasing the individuals into groups of 10, 20 or 50, ensuring that selected pairs were phased together as would be the case in real applications. Reducing the group size reduces phasing accuracy.

#### 3.2.4 New modelling options

Finally, we evaluated the two new modelling options (‘Step’ and ‘Interval’) described above. For this purpose, we again used the 18 genotyped TAF individuals, as well as the bison data. First, we applied a model with 50 ancestral layers, matching the last 50 generations of ancestors, and set the rates of coancestry change to all even numbers between 2 and 100. We then ran models with the ’Step’ option, setting the inbreeding rates to a constant value for groups of five successive generations. We compared the resulting inbreeding rates and levels with those obtained using the default options to demonstrate that the function has been implemented correctly. We expected a similar partitioning, but somehow smoothed, with equivalent inbreeding levels. Then, we ran an equivalent model, defining the HBD class using the ’Interval’ option. Comparing this with the ’Step’ option allows us to demonstrate a successful implementation, as both approaches should result in very similar outcomes. Finally, we ran the last model again using the ’RecTable’ option, which reduces computation time by applying small approximations to the recombination rate. Comparing the results with and without the function allows us to verify that this function improves speed without affecting the results. These analyses also illustrate that the new options produce more interpretable results with inbreeding rate values per generation. Finally, we compared the computational performances of the tested models using TAF individuals.

## 4 RESULTS

### 4.1 Computing time

To illustrate how the new algorithm reduces computing times, we compared runtimes with different models on the TAF datasets. Both package versions produced the same results, but the average time per iteration was 7.9 to 83.8 times shorter with the new package version, depending on the number of fitted HBD classes and the number of markers (see Table 1). As expected, the benefits were maximal when larger numbers of HBD classes were fitted, due to the new linear complexity of the algorithm replacing the former quadratic behaviour. Further information on memory usage is provided in Table S2.

**Table 1.** Comparison of computing times (in seconds) with different versions of the package using models with different number of HBD classes (K). These models were run of the feral cattle populations with different genotyping panels including whole-genome sequence (WGS), high-density (HD) and medium-density (MD) arrays.

| SNP Panel | Package version | K | Rates of coancestry change | Time per iteration |  |  | Time per individual |  |  |
| --- | --- | --- | --- | --- | --- | --- | --- | --- | --- |
|  |  |  |  | Min. | Average | Max. | Min. | Average | Max. |
| WGS | 0.3.2.1 | 15 | {2;4;8;... ;32,768} | 17.050 | 17.104 | 17.206 | 19,038.1 | 24,733.1 | 43,610.1 |
| WGS | 0.4.0 | 15 | {2;4;8;... ;32,768} | 1.505 | 1.563 | 1.648 | 1674.1 | 2303.5 | 4030.5 |
| HD | 0.3.2.1 | 13 | {2;4;8;... ;8192} | 0.899 | 0.904 | 0.920 | 705.3 | 1113.6 | 1578.1 |
| HD | 0.4.0 | 13 | {2;4;8;... ;8192} | 0.099 | 0.104 | 0.107 | 79.0 | 124.7 | 178.8 |
| MD | 0.3.2.1 | 10 | {2;4;8;... ;1024} | 0.020 | 0.021 | 0.021 | 12.3 | 16.7 | 28.0 |
| MD | 0.4.0 | 10 | {2;4;8;... ;1024} | 0.003 | 0.003 | 0.003 | 1.5 | 2.1 | 3.4 |
| MD | 0.3.2.1 | 50 | {2;4;6;... ;100} | 0.894 | 0.902 | 0.913 | 5334.2 | 6980.1 | 9343.9 |
| MD | 0.4.0 | 50 | {2;4;6;... ;100} | 0.010 | 0.011 | 0.012 | 69.2 | 92.2 | 138.3 |

### 4.2 IBD analysis and partitioning for pairs of haplotypes: application to a haploid organism

To illustrate the new IBD analysis feature, we estimated the levels of pairwise IBD among the 40 haploid bee drones (780 pairs). First, we fitted a model with a single IBD class to demonstrate its equivalence with hmmIBD (Schaffner et al., 2018), a reference method for haploid organisms. Despite the different constraints and parameter estimation approaches, the IBD levels estimated with both approaches were highly correlated (r = 0.996), as were the estimated rate of coancestry change parameters (r = 0.984), confirming that the two models are highly similar. We then used a 14-layer model to demonstrate its advantages over a single-class model. The total IBD levels obtained using single-class and multiple-class models were highly correlated (r = 0.982), but the average levels were slightly lower using a single class (0.220 vs. 0.250). With a single-class model, there was no distinction between recent and ancient levels. The estimated rates with this single-class model ranged from 42 to 2340, making them difficult to interpret. Conversely, multiple HBD class models facilitate interpretation through IBD partitioning (see Figure 2 for the distribution of IBD across classes for a random subset of 165 pairs of individuals, and Figure S1 for all the pairs). For example, a concentration of recent IBD relationships is observed in classes with rates of coancestry change of 16 and 32 (average levels of 0.034 and 0.029), consistent with a recent reduction in N_e_ (as reported in Leroy et al. (2024)). There is also a high contribution (0.116) from background IBD levels associated with very short segments (R_c_ = 4,096). Levels associated with more recent demographic events (R_c_ ≤ 32) average at 0.094. Five pairs of individuals have particularly high IBD levels (> 0.45) and clearly stand out. Their IBD partitioning indicates very recent levels (including levels > 0.30 in classes R_c_ = 2 and 4) and shows that the contribution of recent IBD is around 0.50 for these pairs (from 0.380 to 0.615). For these five pairs, the estimated rates of coancestry change using a single-class model ranged from 42 to 126. While these are the five lowest estimated rates, they do not clearly indicate the closeness of the relationships as well as the multiple-class model does.

**Figure 2.**
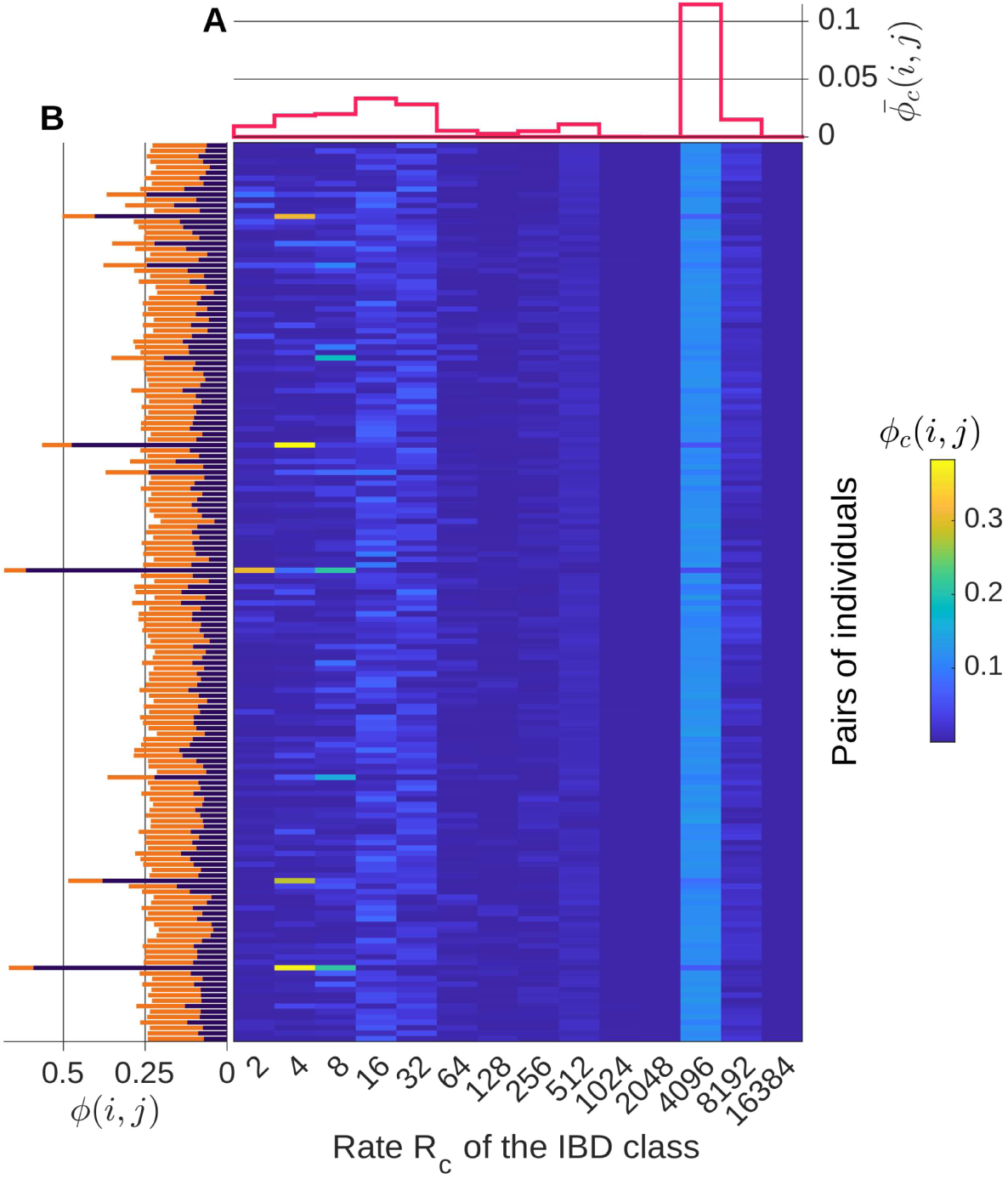
Estimation of IBD sharing for a random subset of 165 pairs of haploid bee drones from the Ouessant Island (out of the 780 pairs). The model estimates the probability, 𝜙(𝑖, 𝑗), that a pair of haplotypes, *i* and *j*, are IBD, and partitions this probability into 14 IBD classes. Within each IBD class c (where c is the index of the class), the length of IBD segments is exponentially distributed with a rate R_c_ (i.e. different IBD classes correspond to different lengths). The central heatmap shows the class-specific pairwise IBD probabilities 𝜙_c_(𝑖, 𝑗), i.e. the probability that haplotypes *i* and *j* share an IBD segment associated with IBD class c. The results are shown for the 165 selected pairs (rows) and the 14 IBD classes (columns), with the color intensity indicating the probability. The two side panels are representing marginal distributions. (**A**) The average IBD probability per class (averaged over all pairs of individuals), denoted 𝜙^ത^ (𝑖, 𝑗). (**B**) Total IBD level for each pair 𝜙(𝑖, 𝑗). These values are obtained using either all IBD classes (orange) or only recent IBD classes with rates R_c_ ≤ 32 (purple).

### 4.3 Relatedness estimation

To illustrate the properties of global and local kinship estimation, we ran a nine-layer model on the 18 genotyped TAF individuals, a population that experienced a severe bottleneck 15-20 generations ago (Gautier et al., 2024). The estimated kinship coefficients represent the probability that two alleles randomly sampled in each individual of a pair are IBD. As before, the partitioning into different IBD classes provides additional information. All pairs of individuals have high relatedness levels (ranging from 0.288 to 0.496, and 0.336 on average), with most IBD captured by classes with rates of coancestry change of 16 and 32 (Figure 3A). In fact, most of the IBD is associated with relatively short segments (< 3 Mb) corresponding to the founder events and a background IBD level in the population. The model allows to separate this background IBD levels present in all individuals (captured mainly by the class with R_c_ = 32) from more recent IBD levels due to closer relationships observed for a few pairs of individuals (captured by classes with rates ≤ 8) (see Figure 3A). We could define relatedness with respect to different base populations, for example by including all classes (and the founder events), or by focusing only on recent IBD (using only the three first classes), as shown in Figure 3B. Interestingly, we observe a pair of individuals with high recent kinship levels (around 0.25), which could correspond to either a parent-offspring pair or a fullsib pair.

**Figure 3.**
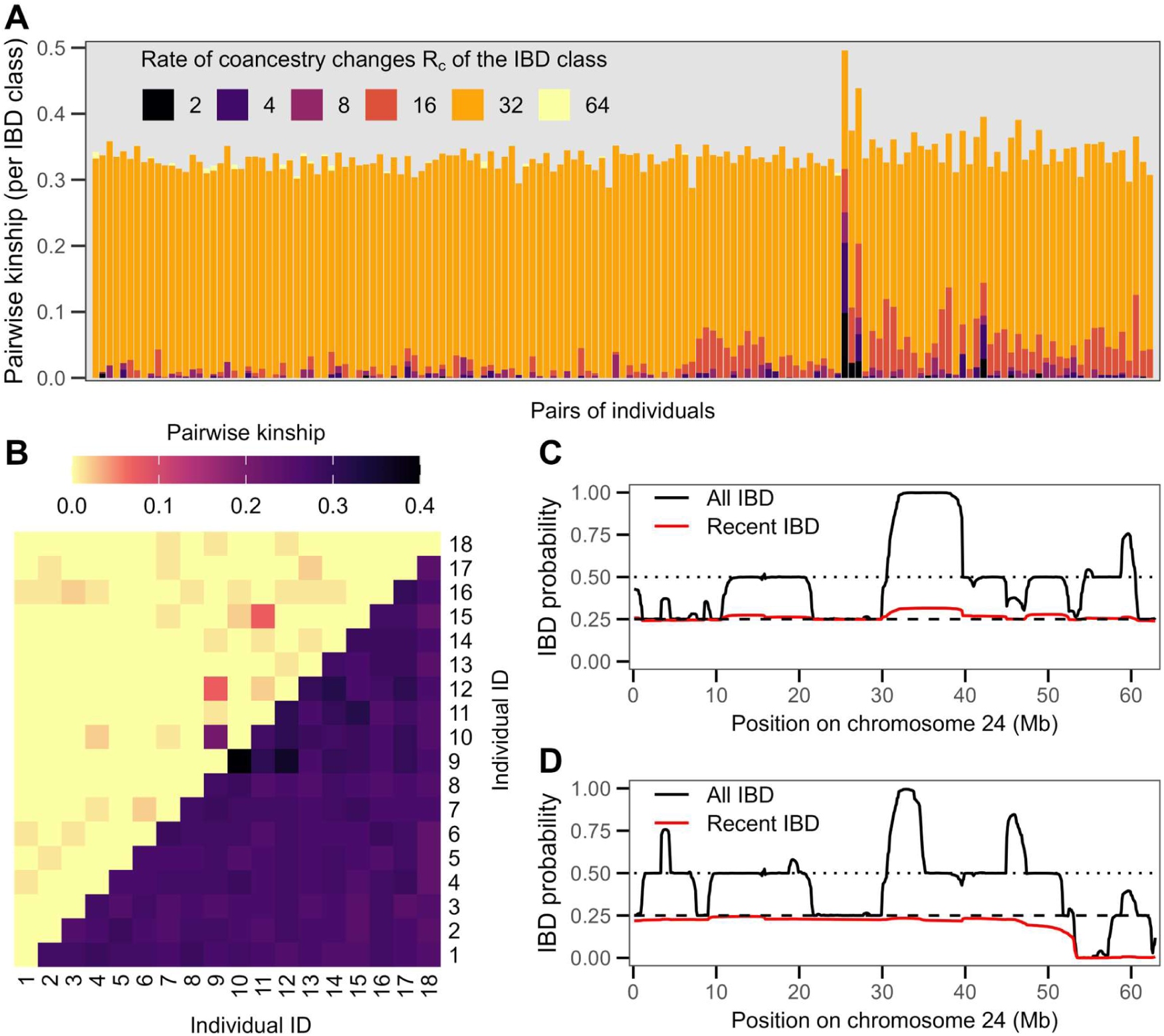
Pairwise kinship estimation among the 18 genotyped TAF individuals. A total of 153 pairs were analyzed. (**A**) Partitioning of the pairwise kinship into different IBD classes defined by their rates of coancestry change R_C_. IBD is mainly associated with the R_C_ = 32 class, and to a lesser extent to the R_C_ = 16 class. Some pairs of individuals also exhibit more recent IBD sharing (R_C_ ≤ 8). HBD classes with R_c_ > 64 are not represented, as their values were null. (**B**) Relatedness matrix among the individuals. Pairwise kinship is estimated using all IBD classes (lower diagonal) or close/recent IBD classes only (upper diagonal). (**C**) Locus-specific IBD probability along chromosome 24 for the pair of individuals with the highest kinship level. This probability is calculated as the probability that two randomly selected haplotypes, one from each individual, are IBD at a specific locus. This value is computed using all IBD classes and only recent/close IBD classes (R_c_ ≤ 8, red). The horizontal dashed lines represent the values corresponding to the sharing of 1 or 2 pairs of IBD haplotypes (at 0.25 and 0.50, respectively). (**D**) Locus-specific IBD probability along chromosome 24 for a second pair of individuals with high kinship levels.

The locus-specific values, corresponding to the probability that two sampled alleles at a locus are IBD, offer additional information for understanding relationships. If there is information to accurately infer the IBD relationships between the four possible pairs of haplotypes, these locus values should take values of 0 (no IBD pair), 0.25 (one pair of IBD haplotypes), 0.50 (two pairs of IBD haplotypes) and 1.00 (the four haplotypes are IBD), and can vary along the chromosomes due to recombination. This is what we observe, for example, for one pair of highly related individuals (Figure 3C). We observe that the locus-specific levels are concentrated at values of 0.25, 0.50 and 1.00, but never 0, suggesting that this is a parent-offspring relationship that should always have at least one pair of IBD haplotypes (and eventually more if both parents are related). This is even clearer when we focus only on recent IBD (R_c_ ≤ 8), as the value is constant at 0.25 along chromosome 24 for example, as expected for this relationship (one pair of IBD haplotypes). The higher levels obtained using all IBD classes indicate that the two parents are related (due to the founder event). Figure 3D shows the local HBD levels for another pair of individuals with some recent relatedness. We observe some regions of null relatedness (no pair of IBD haplotypes), although they remain rare. The number of pairs of IBD haplotypes takes values of 0, 1 or 2, compatible with a full-sib relationship. However, when we use only the most recent relatedness, the value varies from 0 to 1 pair along the genome, more in line with an half-sibs relationship. Please note that the local partitioning between recent and ancient IBD is not always accurate. For example, in the case of the parent-offspring pair, the estimated number of ‘recent’ IBD pairs can be below or above 1 for some genomic positions.

To further evaluate the ability of IBD partitioning to distinguish between recent and background levels of IBD, we ran the same model on pairs of individuals with known relationships in two simulation scenarios. For each pair, we observed recent relatedness associated with long IBD segments (R_c_ ≤ 8), as well as additional relatedness associated with more distant ancestors (R_c_ > 8) (Figure 4). The recent levels matched those expected for different relationships, and were null for unrelated individuals (Figure 4). In this context, we also compared the properties of RZooRoH with those of KING (Manichaikul et al., 2010). KING is a reference method for estimating relatedness from genotyping data, proposing two complementary approaches that can be eventually combined: a robust kinship estimator and one based on IBD segments. For each pair category, the estimators’ distributions were centred around the expected value for both RZooRoH and the KING robust kinship estimator. The best estimator varied depending on the relatedness category and scenario. In general, KING estimators exhibited greater variability, and could also take negative values. Identifying unrelated individuals was quite efficient with RZooRoH, as these pairs did not have any recent IBD segments. The KING estimator based on IBD segments was systematically higher than the expected levels because it also captures background inbreeding (see Figure S2). The same was true when all IBD segments were used with RZooRoH, although RZooRoH captured even higher background levels.

**Figure 4.**
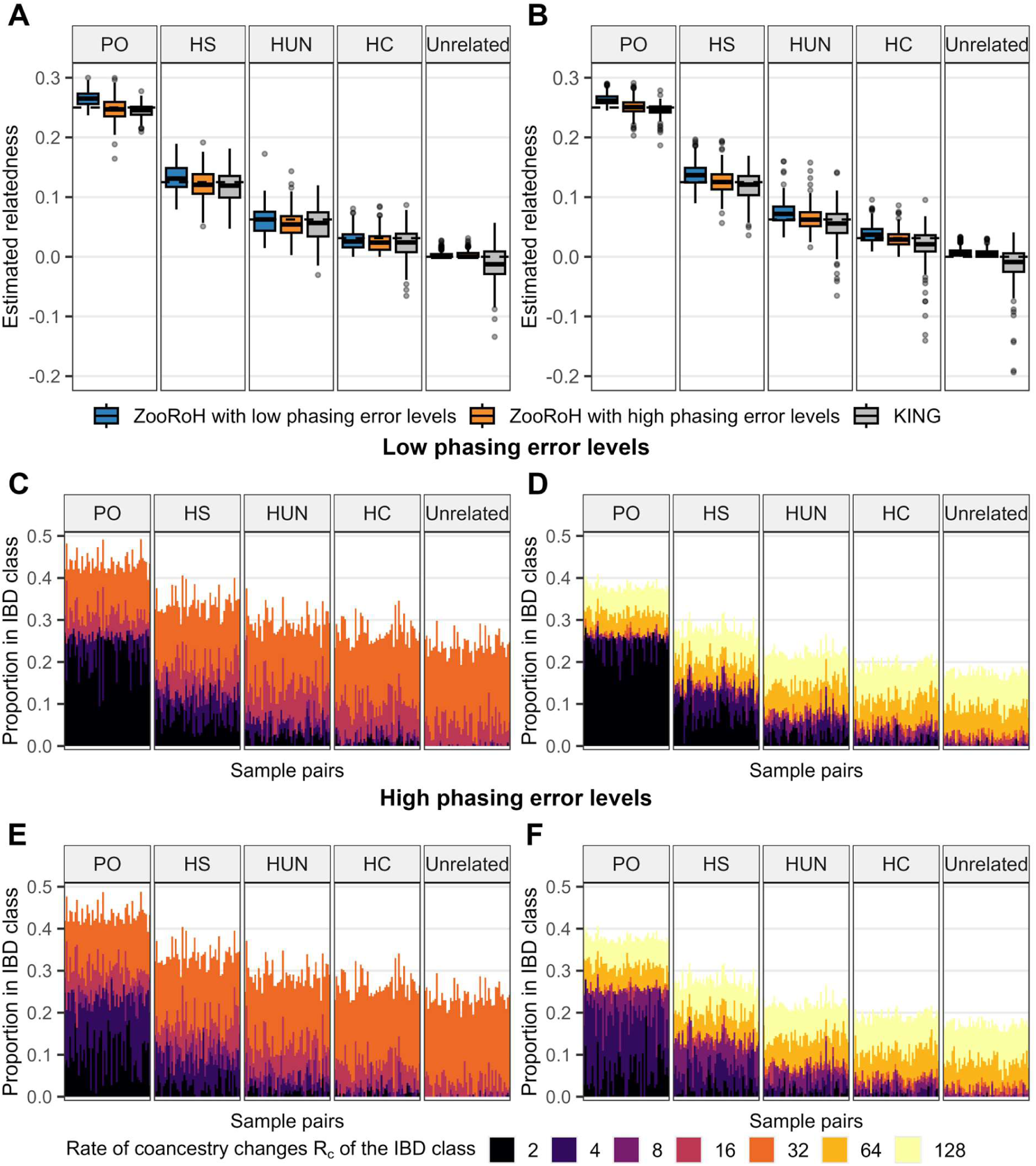
Estimation of pairwise kinship on simulated data sets. (**A**) Boxplots of estimated relatedness using RZooRoH and with recent IBD only (R_c_ ≤ 8), and two levels of phasing error (low and high phasing errors correspond to phasing all individuals jointly or in subsets of 20), and with the KING robust estimator. The relatedness is reported for the following kinship categories: parent-offspring (PO), half-siblings (HS), half-uncle-nephew (HUN), half-cousins (HC), and unrelated individuals (no common ancestor in the last three generations). The horizontal dashed lines indicate the expected kinship level for each relationship. Results are reported for the scenario with a bottleneck. (**B**) Same as panel A, but for the second scenario with a constant N_e_. (**C**) Partitioning of the pairwise kinship into different IBD classes defined by their rates R_C_ for the different categories of kinship and the first scenario. Phasing was done jointly for all individuals (i.e. low phasing error level). (**D**) Same results but for the second simulation scenario. (**E**,**F**) Same as panels **C** and **D**, but with phasing done in subsets of 20 individuals (i.e. high phasing error level).

The analysis was then repeated to study the impact of phasing errors generated by phasing individuals into smaller groups. Switch error rates (SER) increased when individuals were phased in smaller groups (Table S3). In the first scenario, SER were around 1-2% when phasing was performed on groups of 20 individuals and above 5% for groups of 10. Even larger SER were observed for the second scenario. These SER are extremely high, particularly when we consider that, for IBD segment identification, errors can occur in both haplotypes. These high SER enable us to study the impact of these errors on relatedness estimation. As the SER increased, the level of recent inbreeding captured tended to decrease slightly, while total inbreeding remained unchanged (Figures S2 & S3). When phasing was performed on subsets of 20 or 50 individuals, the distinction between recent and background inbreeding remained effective (Figure S3), although a shift towards slightly longer IBD segments was observed. For example, recent IBD was captured in classes with R_c_ ≤ 4 when there were few phasing errors. With higher SER, however, some IBD segments moved to the R_c_ = 8 class, and this effect became more pronounced as the SER increased (see Figure 4D & S4). When phasing subset of 10 individuals, IBD segments were divided further and IBD shifted towards classes with higher rates of coancestry change (Figure 4 & S4). Consequently, the estimated kinship levels using classes with R_c_ ≤ 8 were lower than expected. Overall, these analyses demonstrate that, by partitioning IBD into different length-based classes, it is possible to distinguish between recent and ancient inbreeding. However, phasing errors may require redefining the thresholds used to capture recent IBD. Indeed, these errors tend to divide IBD segments into shorter ones, shifting the distribution towards shorter categories, although this has less of an impact on more ancient IBD (see Figure S2).

### 4.4 New modelling options

To illustrate the partitioning of HBD using the new modelling options, as well as their properties, we used the cattle (TAF) and European bison datasets. These options were designed to refine modelling for more advanced use, offering more interpretable results by reporting inbreeding rates per generation and providing a clearer definition of the boundaries of HBD classes. We began by fitting a model with 50 HBD classes with their respective rates of coancestry changes matching those from ancestors present in the last 50 generations (one class per generation). These two populations experienced a recent severe bottleneck ca. 15-20 and 10-15 generations ago (see Gautier et al. (2024) and Druet et al. (2020) for details), and high HBD levels were observed, concentrated in recent HBD classes (Figure 5). For example, in the 18 genotyped TAF, HBD was concentrated in five classes with rates comprised between 26 and 34 (corresponding to a bottleneck approximately 15 generations ago). When HBD proportions were plotted per individual (Figure S5), we observed that they were concentrated in even fewer classes, only one or two per individual. The same behaviour was observed for European bison (Figure S6), where HBD was concentrated in more recent classes (the average levels maximizing in classes with rates between 8 and 24).

**Figure 5.**
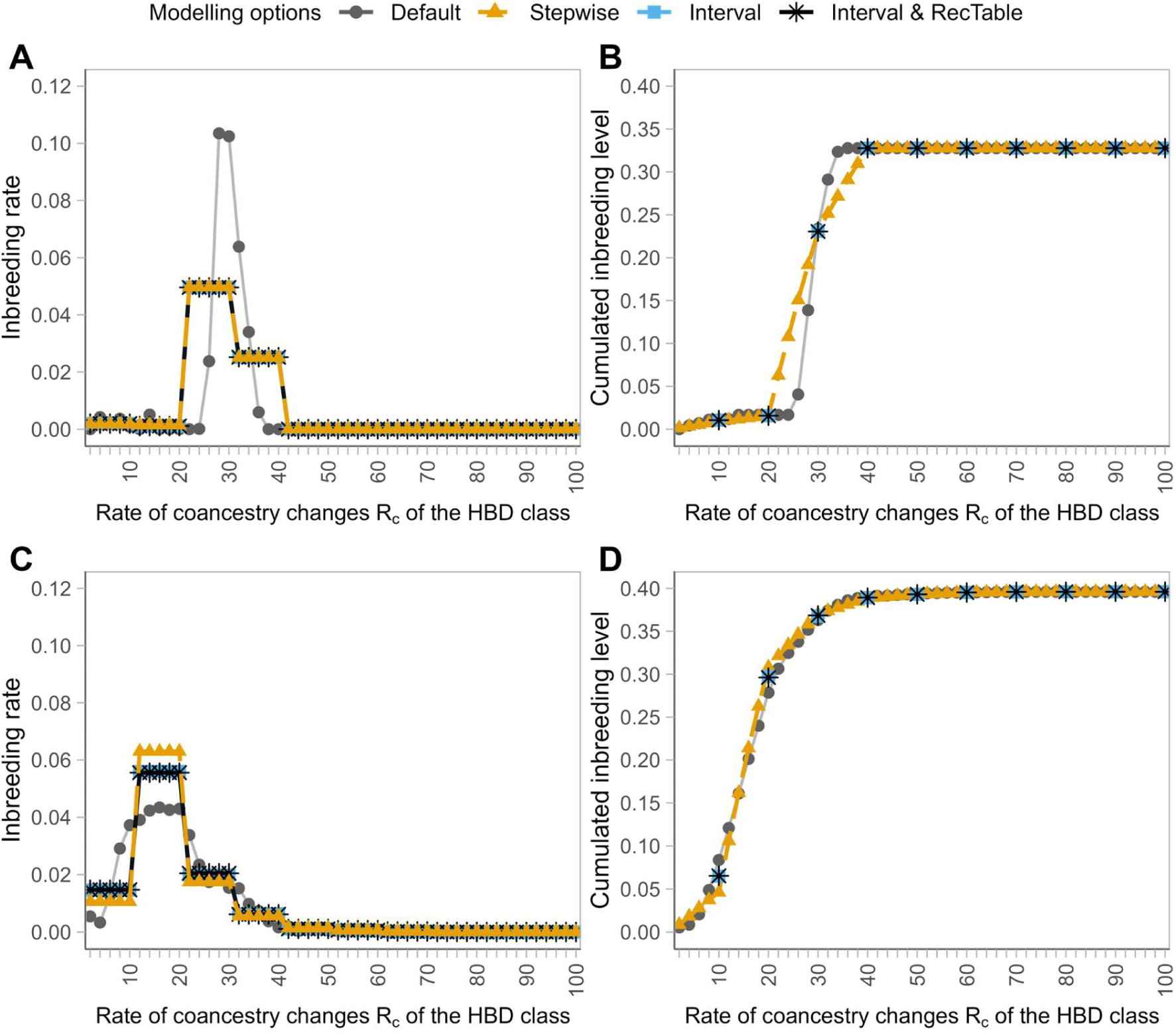
Estimation and partitioning of HBD levels with new modelling options. Estimation was performed using a model with 50 layers and default options, with rates of coancestry change ranging from 2 to 100 in steps of 2 (approximately one layer per generation of ancestors). Next, the ‘Step’ option was used to impose that the inbreeding rates per layer remained constant for ten groups of five classes (2-10, 12-20, etc). Finally, a similar model was fitted with the ‘Interval’ option, with HBD classes defined as 10 intervals of 5 layers (corresponding to generations 1 to 5, 6 to 10, etc). The ‘Interval’ model was run with or without the RecTable option, which reduces computation costs by making certain approximations. (**A**) Estimated inbreeding rates per generation in the TAF cattle population. (**B**) Cumulative realized inbreeding levels in the TAF population when an increasing number of HBD classes are used. (**C**) Estimated inbreeding rates per generation in the European bison population. (**D**) Cumulative realized inbreeding levels in the European bison population when an increasing number of HBD classes are used.

The overall trends were similar when using the ‘Step’ option (with a constant inbreeding rate for groups of five consecutive HBD classes), although the HBD was consequently distributed over more layers in the TAF (Figure 5). With the ‘Interval’ option, the results were very similar to those obtained with the ‘Step’ option (Figure 5), although small differences were observed in the European bison.

The overall inbreeding coefficients were identical with the different approaches (Figure S7). The comparison of the curves showing the cumulated inbreeding levels at different points in the past (Figure 5) shows that all methods capture the same inbreeding levels, but with slightly different partitioning. The comparison of the results obtained using the ’Step’ and ’Interval’ options shows that these models are very similar. The former still fits one HBD class per generation, limiting its range, while the latter fits one HBD class per group of generations through more complex computations of transition probabilities. Finally, we tested whether the ‘RecTable’ option, which approximates the genetic map to reduce computation time, provided similar results. The analyses with and without the option were hardly distinguishable (Figure 5), and the distributions of HBD segments were almost identical (Figure S8). Compared to the default model, using these new options allows us to refine the timing of the event and provide more interpretable inbreeding rates. For example, values between 0.05 and 0.10 would correspond to N_e_ values of between 5 and 10 individuals at some point in the past, which is consistent with historical records.

The computational efficiency of these different approaches is reported in Tables S2 and S4. The number of iterations required for parameter estimation is significantly reduced when the ‘Step’ option is used (approximately tenfold with groups of five HBD classes). This indicates that faster convergence is achieved when fewer parameters need to be estimated. In general, when testing the package on various datasets, we observed that models with fewer parameters experienced fewer convergence failures (i.e. optim reported an error status or reached the iteration limits). With the ‘Interval’ option, the number of iterations is about the same as with equivalent ‘Step’ models, but the running times are about twice as long. Using the ‘RecTable’ option allows a tenfold reduction in computing time, resulting in very short running times, although the model with groups of five HBD classes is not faster than a classical model with 10 layers. Note that the ‘Step’ approach is limited to low or medium density arrays because at higher densities, more than 1,000 past generations can be captured, and this would require fitting too many HBD classes. We tested whether it was possible to run an ‘Interval’ model that captured more past generations of ancestors at the MD, HD or WGS levels, both with and without the ‘RecTable’ option (Table S4). Without the ‘RecTable’ option, this approach was inefficient. For example, individual estimations did not converge within the 48-hour time limit when WGS data was used. However, using the ‘RecTable’ option enabled us to run models within a more reasonable time: an average of 0.23, 5.33 and 54.65 minutes, respectively, according to the marker panel.

## 5. DISCUSSION

Here, we present substantial improvements to the RZooRoH package, including a significant reduction in running times, an extension to the analysis of haploid individuals and the estimation of relatedness, and new modelling options. The reduction of running times is a key feature, particularly as it makes the other applications possible.

Thanks to the new formats for reading haploid and phased data, the package can now be used to study haploid organisms such as bee drones (Leroy et al., 2024), *Plasmodium falciparum* (Schaffner et al., 2018) and *Chlamydomonas reinhardtii* (Craig et al., 2019), for which IBD analyses were previously performed using a similar HMM called hmmIBD (Schaffner et al., 2018). As shown in the results section, this HMM is highly similar to the ZooRoH model with a single IBD class. Using multiple IBD classes provides more information about kinship and recent demographic history, as demonstrated using the honey bee dataset. Similarly, it can be applied to the sex-chromosomes in the heterogametic sex. This technique is sometimes used in humans to study past demographic events without introducing phasing errors (e.g., Fernandes et al., 2021).

The ability to model IBD between pairs of chromosomes from different individuals makes it possible to model the relationship between two individuals, since this depends on the IBD relationship between the four possible pairs of haplotypes from these two individuals (with each individual contributing one haplotype to each pair). Their kinship can be estimated as the probability that one of the four possible pairs is IBD, and this approach can provide both global and local estimators of realized relatedness. We have previously demonstrated the efficiency of the ZooRoH model for such applications, including estimating relatedness between two individuals and predicting inbreeding levels in the (future) offspring of genotyped parents (Forneris et al., 2025). Therefore, we have integrated it into RZooRoH to make it easier for interested users to apply. Predicting future HBD levels could be useful for mating plans to manage the genetic diversity of populations (e.g., de Cara et al., 2013). As illustrated in the present study, the ability to distinguish between recent and ancient segments, the latter of which results from demographic events affecting the entire population, helps to infer the relationship between two individuals. In the feral cattle population, for instance, we observed that all individuals had high relatedness associated with segments of moderate length (around 3 Mb). However, some pairs of individuals also shared longer IBD segments, and corresponded to parent-offspring (sharing one IBD haplotype across the entire genome) and half-sibling relationships (sharing 0 or 1 IBD haplotype across the genome), respectively.

Analyses of simulated data confirmed that the model can distinguish recent from more ancient relatedness. However, estimating relatedness in diploid individuals requires a phasing step. This step is assumed not to introduce too many errors, an assumption made by many models designed to identify IBD segments (e.g., Freyman et al., 2021; Zhou et al., 2020) or infer demographic history from IBD segments (e.g., Fournier et al., 2023). Recent studies have shown that statistical phasing can be achieved accurately (e.g., Oget-Ebrad et al., 2022), and IBD estimation can be accurate when phased haplotypes are used (e.g., Forneris et al., 2025). To further study the impact of phasing errors, we applied the model to datasets with increasing levels of phasing errors and observed that performances can deteriorate (e.g., distinguishing between recent and background relatedness becomes more difficult). Generally, long IBD segments were divided into shorter ones, with IBD shifting towards IBD classes with higher rates of coancestry change (i.e. shorter lengths). However, phasing errors had less impact on relatedness levels measured with respect to more distant base populations.

To help users understand the benefits of this new relatedness estimator, Table S5 compares its properties with those of other popular approaches. Like KING, RZooRoH can distinguish between recent and background relatedness, although it relies on different information (the length of IBD segments), which makes the two approaches complementary. Both approaches also provide information on total IBD levels, as measured by all IBD segments. One difference is that RZooRoH provides a finer classification, as IBD is partitioned into multiple classes, whereas KING provides two extreme values: robust kinship estimators (corrected for sample composition and structure), and estimators based on all identified IBD segments without correction. Many other estimators behave either as the robust kinship estimator, providing a single estimate corrected for population levels, or as the IBD-based estimator, providing an estimate based on all identified IBD segments. In addition to the aforementioned applications, RZooRoH’s main properties include its reliance on IBD segments, which provides additional information; its range always being between 0 and 1, unlike many SNP-based approaches; and its provision of locus-specific information, as well as information on the number of pairs of haplotypes that are IBD (information that can be used to further characterize relationships). However, it is not computationally efficient compared to other methods and should not be used for large samples or applications involving phenotypes, such as genomic selection, variance components estimation or genome-wide association studies. Unlike methods such as KING or other SNP-based estimators of genomic relationships, the approach is also sensitive to high levels phasing errors when used for applications focusing on the most recent IBD classes.

Finally, the new modelling options provide tools for more refined analyses, offering a clearer definition of the boundaries between HBD classes and a finer partitioning resolution. They also provide estimates of inbreeding rates per generation. However, the (default) layer model remains faster while still providing estimates of inbreeding coefficients, performing HBD partitioning and providing qualitative information about the number of generations to common ancestors.

Overall, the new version of RZooRoH brings several important enhancements. Some of these, such as relatedness estimation and the use of ’Interval’ models to estimate inbreeding rates, require further evaluation. We are therefore currently conducting such evaluations to better characterize their properties. These new features and other minor changes are described in detail in the updated vignette. We anticipate the new version of the package will be useful to a broad range of researchers.

## ACKNOWLEDGEMENTS

We acknowledge Thibault Leroy for sharing the original honey bee whole-genome sequencing data set. We would like to thank Samuel Speak and the three anonymous reviewers whose valuable comments helped us to improve the manuscript. Tom Druet is Research Director from the Fonds de la Recherche Scientifique—FNRS (F.R.S-FNRS). Computations were carried out using the supercomputing facilities of the “Consortium d’Equipements en Calcul Intensif en Fédération Wallonie-Bruxelles” (CECI), funded by the F.R.S-FNRS. This work was supported by Fonds De La Recherche Scientifique—FNRS, ROAGE T.0102.24. Service Public de Wallonie, BEWARE FitSel project—convention no. 2110192.

## DATA ACCESSIBILITY STATEMENT

The real data sets used in this study were previously generated and made publicly available by other contributors. All SNP genotyping data for the TAF individuals are publicly available from the WIDDE repository (http://widde.toulouse.inra.fr/widde/). WGS data for the 8 TAF individuals are available from the NCBI SRA repository (see Supplementary Table S3, Supplementary Material online in Gautier et al. (2024) for accession run IDs). The genotyping data for the individuals from the European bison population are available on Dryad (doi:10.5061/dryad.xsj3tx9b7). All sequencing data from the honey bee dataset are publicly accessible, with corresponding SRA accession numbers provided in supplementary table S1 and supplementary File S1, Supplementary Material online in Leroy et al. (2024).

## Notes

### Competing Interest Statement

The authors have declared no competing interest.

### Summary of Updates

This version contains more accessible explanations (more user-oriented) of the updates of the package and additional analyses to illustrate new features (for example, for the estimation of relatedness). Supplemental files have been also updated.

