## Supplementary Material for "Updating the RZooRoH package for the analysis of inbreeding, identity-by-descent and relatedness from genomic data"

Table S1. Description of simulated datasets. Simulations were performed with SLiM 4.0.1 [1] and msprime 1.2.0 [2] using forward-time simulation and a “recapitation” technique [3].

| Parameter | Population with bottleneck | Constant size population |
|---|---|---|
| Initial population | 10,000 diploid individuals | 10,000 diploid individuals |
| Demographic evolution | $N_e$ = {1000, 500, 200, 50, 10, 1000} for generation intervals = {1-4; 5-8; 9-12; 13-15; 16-20; 21-1000} | $N_e$ = 250 for generations 1 to 1000 |
| Genomes size | 25 chromosomes pairs of 100 cM each | 25 chromosomes pairs of 100 cM each |
| Recombination rate | 1e-8 per bp | 1e-8 per bp |
| Mutation rate | 1e-8 per bp | 1e-8 per bp |
| Mating | Random | Random |
| Sex ratio | 1 | 1 |
| Marker selection | 25,000 evenly spaced bi-allelic markers with a MAF ≥ 0.01 | 25,000 evenly spaced bi-allelic markers with a MAF ≥ 0.01 |
| Marker density | 10 SNPs / cM | 10 SNPs / cM |

Table S2. Memory requirements (in kilobytes) with different versions of the package using models with different number of HBD classes (K) and options. These models were run of the feral cattle populations with different genotyping panels including whole-genome sequence (WGS), high-density (HD) and medium-density (MD) arrays.

| SNP panel | Package version | K | Rates of coancestry changes | Options | Max Resident Set Size |
|---|---|---|---|---|---|
| WGS | 0.3.2.1 | 15 | {2;4;8;… ;32,768} | Default | 9361136K |
| WGS | 0.4.0 | 15 | {2;4;8;… ;32,768} | Default | 9945632K |
| HD | 0.3.2.1 | 13 | {2;4;8;… ;8192} | Default | 794360K |
| HD | 0.4.0 | 13 | {2;4;8;… ;8192} | Default | 891120K |
| MD | 0.3.2.1 | 10 | {2;4;8;… ;1024} | Default | 129528K |
| MD | 0.4.0 | 10 | {2;4;8;… ;1024} | Default | 131300K |
| MD | 0.3.2.1 | 50 | {2;4;6;… ;100} | Default | 157828K |
| MD | 0.4.0 | 50 | {2;4;6;… ;100} | Default | 166044K |
| WGS | 0.4.0 | 15 | {2-4;6-8;10-16;…;32,770-65,536} | Interval & RecTable | 10992484K |
| HD | 0.4.0 | 13 | {2-4;6-8;10-16;…;8194-16,384} | Interval & RecTable | 968972K |
| MD | 0.4.0 | 50 | {2;4;6;… ;100} | Default | 166044K |
| MD | 0.4.0 | 50 | {2;4;6;… ;100} | Step 5† | 168908K |
| MD | 0.4.0 | 50 | {2;4;6;… ;100} | Step 10† | 186940K |
| MD | 0.4.0 | 10 | {2-10;12-20;22-30;…;92-100} | Interval | 131164K |
| MD | 0.4.0 | 10 | {2-10;12-20;22-30;…;92-100} | Interval & RecTable | 133464K |
| MD | 0.4.0 | 5 | {2-20;22-40;42:60;…;82-100} | Interval | 129400K |
| MD | 0.4.0 | 5 | {2-20;22-40;42:60; …;82-100} | Interval & RecTable | 129896K |
| MD | 0.4.0 | 10 | {2-4;6-8;10-16;…;1026-2048} | Interval | 130988K |
| MD | 0.4.0 | 10 | {2-4;6-8;10-16;…;1026-2048} | Interval & RecTable | 131308K |

†The value indicates number of layers grouped with the 'Step' option

Table S3. Switch error rates in the two simulated scenarios. Switch error rates are counted per heterozygous marker, and the number of errors per Mb are also reported. Errors are reported as a function of relatedness and the number of individuals in the phasing group. 'All' indicates that the entire last three generations were phased together (3,000 individuals in scenario 1 and 750 individuals in scenario 2).

| Relationship | Phasing group size | Scenario 1 | | Scenario 2 | |
|---|---|---|---|---|---|
| | | SER | Switches per Mb | SER | Switches per Mb |
| Parent-offspring | All | 0.0010 | 0.003 | 0.0008 | 0.002 |
| | 50 | 0.0035 | 0.012 | 0.0065 | 0.018 |
| | 20 | 0.0105 | 0.036 | 0.0308 | 0.084 |
| | 10 | 0.0544 | 0.184 | 0.0972 | 0.266 |
| Half-siblings | All | 0.0011 | 0.004 | 0.0009 | 0.002 |
| | 50 | 0.0060 | 0.020 | 0.0109 | 0.030 |
| | 20 | 0.0156 | 0.053 | 0.0455 | 0.125 |
| | 10 | 0.0645 | 0.218 | 0.1227 | 0.336 |
| Half-uncle-nephew | All | 0.0010 | 0.003 | 0.0008 | 0.002 |
| | 50 | 0.0070 | 0.024 | 0.0125 | 0.034 |
| | 20 | 0.0179 | 0.061 | 0.0516 | 0.141 |
| | 10 | 0.0693 | 0.235 | 0.1335 | 0.366 |
| Half-cousins | All | 0.0011 | 0.004 | 0.0009 | 0.002 |
| | 50 | 0.0078 | 0.026 | 0.0139 | 0.038 |
| | 20 | 0.0197 | 0.067 | 0.0566 | 0.155 |
| | 10 | 0.0723 | 0.245 | 0.1408 | 0.386 |
| Unrelated | All | 0.0011 | 0.004 | 0.0009 | 0.002 |
| | 50 | 0.0084 | 0.028 | 0.0147 | 0.040 |
| | 20 | 0.0205 | 0.070 | 0.0602 | 0.165 |
| | 10 | 0.0747 | 0.253 | 0.1456 | 0.399 |

Table S4. Computing times with different class definitions and new options. We compared equivalent models defined using either the 'Step' or the 'Interval' options (i.e. the number of fitted generations were identical, as well as the range of the groups with identical inbreeding rate) using the MD panel. In addition, the Interval option was also evaluated on other panels and with models including more past generations.

| SNP panel | K | Rates of HBD classes | Class definition and options | Time per iteration | | | Time per individual | | | Number of iterations | | |
|---|---|---|---|---|---|---|---|---|---|---|---|---|
| | | | | Min. | Average | Max. | Min. | Average | Max. | Min. | Average | Max. |
| MD | 50 | {2;4;8;…;100} | SingleRate | 0.010 | 0.011 | 0.012 | 69.2 | 92.1 | 138.3 | 5757 | 8590.6 | 13635 |
| MD | 50 | {2;4;8;…;100} | SingleRate & Step 5† | 0.010 | 0.012 | 0.014 | 6.9 | 10.2 | 13.8 | 651 | 849.3 | 1029 |
| MD | 10 | {2-10;12-20;22-30;…;92-100} | Interval | 0.023 | 0.023 | 0.026 | 15.4 | 20.3 | 24.3 | 672 | 874.6 | 1050 |
| MD | 10 | {2-10;12-20;22-30;…;92-100} | Interval & RecTable | 0.003 | 0.003 | 0.003 | 2.0 | 2.5 | 3.3 | 693 | 878.5 | 1218 |
| MD | 50 | {2;4;8;…;100} | SingleRate & Step 10† | 0.011 | 0.011 | 0.012 | 3.7 | 4.7 | 5.7 | 330 | 425.9 | 484 |
| MD | 5 | {2-20;22-40;42:60;…;82-100} | Interval | 0.019 | 0.019 | 0.020 | 5.6 | 8.3 | 13.7 | 286 | 427.8 | 715 |
| MD | 5 | {2-20;22-40;42:60; …;82-100} | Interval & RecTable | 0.002 | 0.002 | 0.002 | 0.6 | 0.8 | 1.4 | 286 | 418.6 | 693 |
| MD | 10 | {2-4;6-8;10-16;…;1026-2048} | Interval | 0.356 | 0.357 | 0.358 | 247.5 | 392.8 | 599.3 | 693 | 1101.3 | 1680 |
| MD | 10 | {2-4;6-8;10-16;…;1026-2048} | Interval & RecTable | 0.013 | 0.013 | 0.013 | 9.02 | 14.3 | 21.0 | 714 | 1131.7 | 1680 |
| HD | 13 | {2-4;6-8;10-16;…;8194-16,384} | Interval | more than 24h‡ | | | | | | | | |
| HD | 13 | {2-4;6-8;10-16;…;8194-16,384} | Interval & RecTable | 0.194 | 0.199 | 0.201 | 292.2 | 320.2 | 345.3 | 1458 | 1613.2 | 1782 |
| WGS | 15 | {2-4;6-8;10-16;…;32,770-65,536} | Interval | more than 48h‡ | | | | | | | | |
| WGS | 15 | {2-4;6-8;10-16;…;32,770-65,536} | Interval & RecTable | 2.075 | 2.098 | 2.118 | 2251.0 | 3278.6 | 4732.0 | 1085 | 1561.6 | 2263 |

†The value indicates number of layers grouped with the 'Step' option; ‡The running time exceeded the selected time limit for the jobs

Table S5. Properties of ZooRoH relatedness estimation in comparison with other common approaches. For each of the approaches, we used one (or two) representative software to illustrate the properties, some of these may change with other software. The general properties of the different methods are indicative and have not been evaluated in the present study.

| Approach | RZooRoH | Robust moment-estimator | Genomic Relationship Matrix | IBD segments identification | Modelling IBD along chromosomes |
|---|---|---|---|---|---|
| **Software example** | | KING[1] [4] | GCTA [5] | Phaseibd [6], IBDkin [7] | localNGSrelate [8], IBD_Haplo [9] |
| **Input** | Phased genotypes | Genotypes | Genotypes | Phased genotypes | Both[2] |
| **Output** | Kinship coefficient, local probabilities & IBD segments | Kinship coefficient | Genomic Relationship Matrix | Kinship coefficient & IBD segments | Kinship coefficient & local probabilities |
| **Large data sets / speed** | + | ++++ | ++++ | +++ | ++ |
| **Background level estimation[3]** | Yes | No | No | Yes[4] | Yes[4] |
| **Background level correction[5]** | Yes (by class selection) | Yes | Yes | No[4] | No[4] |
| **Negative estimates** | No | Yes | Yes | Typically no | No |
| **Locus-specific values** | Yes | No | No | Needs post-processing | Yes |
| **IBD configuration[6]** | Global and local | Global | No | Global and local | Global and local |
| **Partitioning of IBD** | Yes | No | No | No | No |
| **Typical applications** | Prediction of future inbreeding (global and local), relatedness in small populations, separating recent from background kinship[7], understanding kinship | GWAS, detecting family relationships, selecting unrelated individuals, relatedness in small populations | Genomic prediction, variance components estimation, GWAS, polygenic traits analysis | Using IBD segment to detect and refine close relationships in genetic genealogy services, targeted applications: positive selection, disease mapping, parent-of-origin effects, etc. | Applications with low-coverage sequencing data and with low-confidence genotypes, ancient DNA, small populations of non-model organism, more distant relationships, disease mapping |

[1]We refer to the robust estimator (KING has also an estimator based on IBD segments; [2]These approaches can also use genotype likelihoods; [3]This row indicates whether the estimator provides information on background levels; [4]These approaches report all captured IBD / IBD segments – the segment length limit determines how far relatedness is captured; [5]This row indicates whether the kinship estimators are corrected for background levels in the population, or expressed relative to the population levels; [6]Indicates whether the approach provides individual information useful to classify the relationship, such as information on IBD sharing states (0, 1 or 2 shared haplotypes), the Jacquard's identity coefficients, etc.; [7]Thanks to the different IBD classes, users have more flexibility in determining how to make the separation.

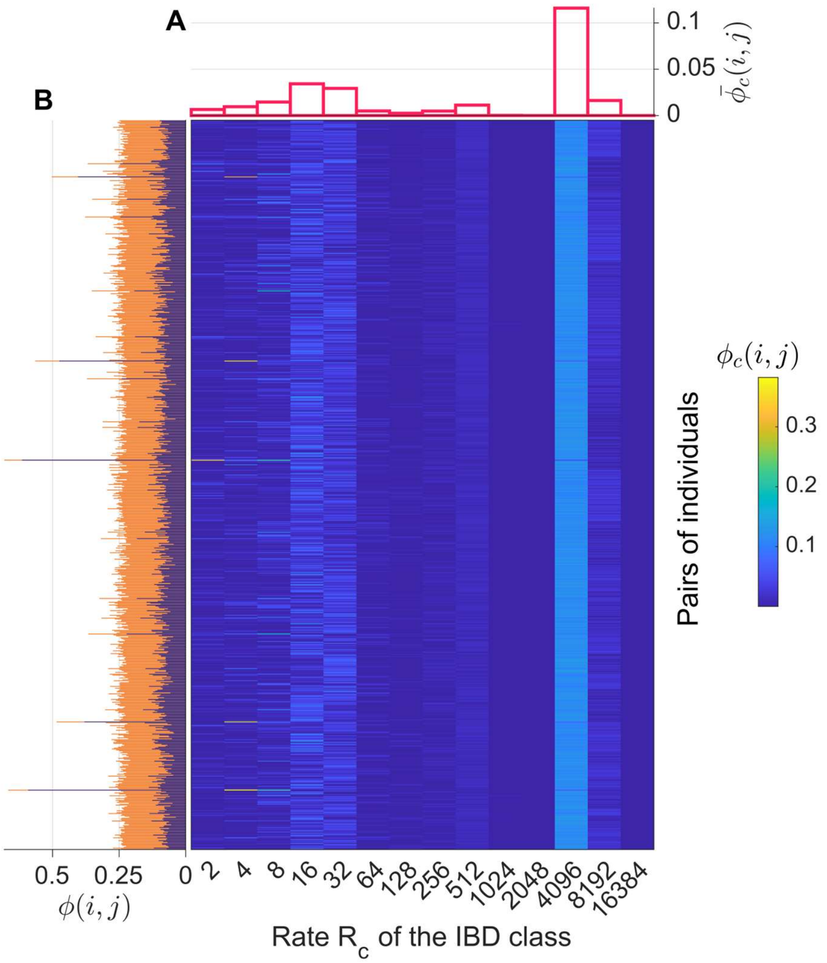

Figure S1. Estimation of IBD sharing for the 780 pairs of haploid bee drones from the Ouessant Island. The model estimates the probability, $\phi(i,j)$, that a pair of haplotypes, *i* and *j*, are IBD, and partitions this probability into 14 IBD classes. Within each IBD class c (where c is the index of the class), the length of IBD segments is exponentially distributed with a rate $R_c$ (i.e. different IBD classes correspond to different lengths). The central heatmap shows the class-specific pairwise IBD probabilities $\phi_c(i,j)$, i.e. the probability that haplotypes *i* and *j* share an IBD segment associated with IBD class c. The results are shown for the 780 pairs (rows) and the 14 IBD classes (columns), with the color intensity indicating the probability. The two side panels are representing marginal distributions. (**A**) The average IBD probability per class (averaged over all pairs of individuals), denoted $\bar{\phi}_c(i,j)$. (**B**) Total IBD level for each pair $\phi(i,j)$. These values are obtained using either all IBD classes (orange) or only recent IBD classes with rates $R_c \leq 32$ (purple).

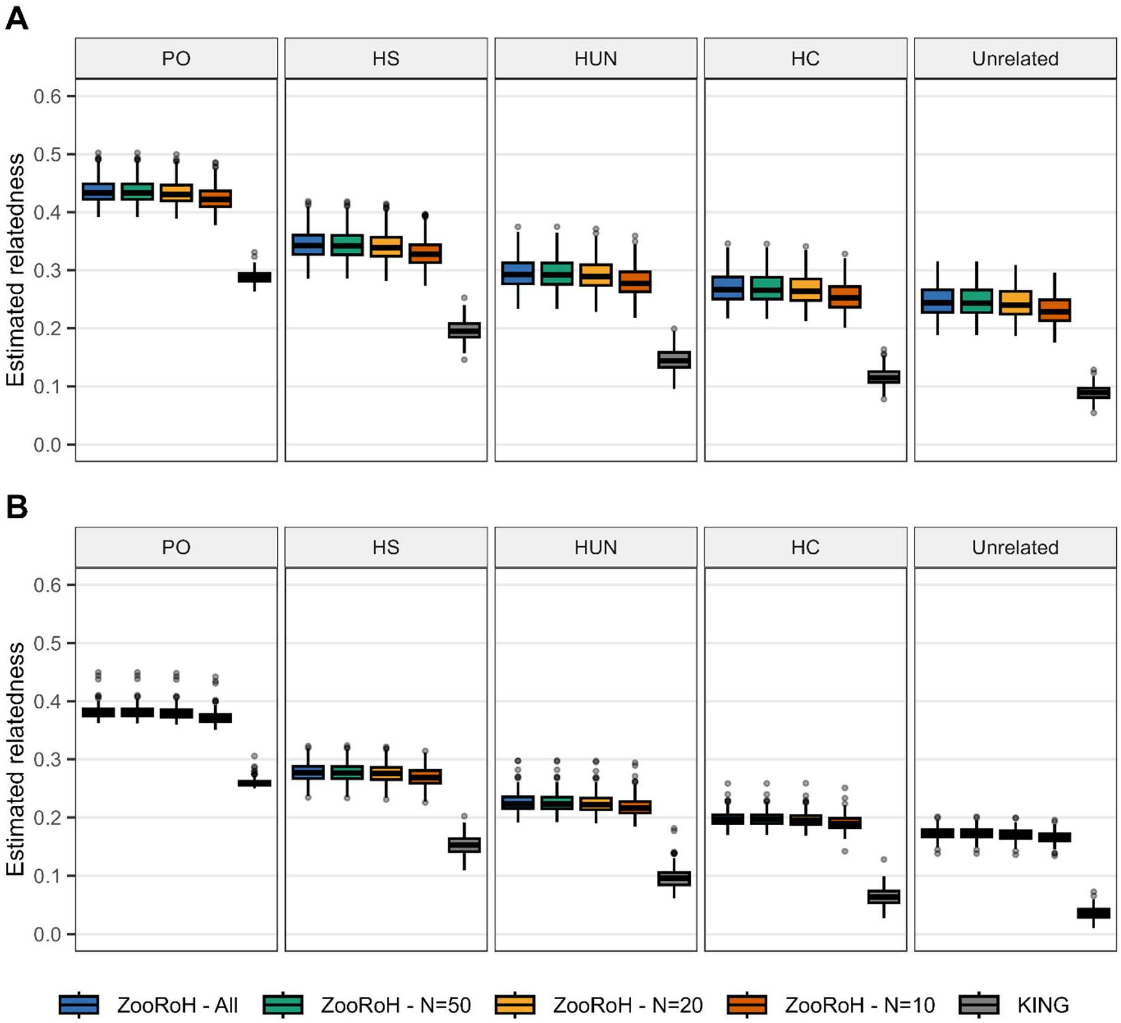

Figure S2. Estimation of pairwise kinship using all IBD classes on simulated data sets. Boxplots showing estimated relatedness with ZooRoH using all IBD classes, with four levels of phasing error, and with the KING IBD estimator. The levels of phasing errors are indicated by the number of individuals phased per group: ALL, N=50, N=20 and N=10 individuals, which correspond to low, moderate, high and very high error levels, respectively. Relatedness is reported for the following kinship categories: parent–offspring (PO), half-siblings (HS), half-uncle–nephew (HUN), half-cousins (HC) and unrelated individuals (no common ancestor in the last three generations). (**A**) Results for the first scenario, with a bottleneck. (**B**) Results for the second scenario, with a constant $N_e$.

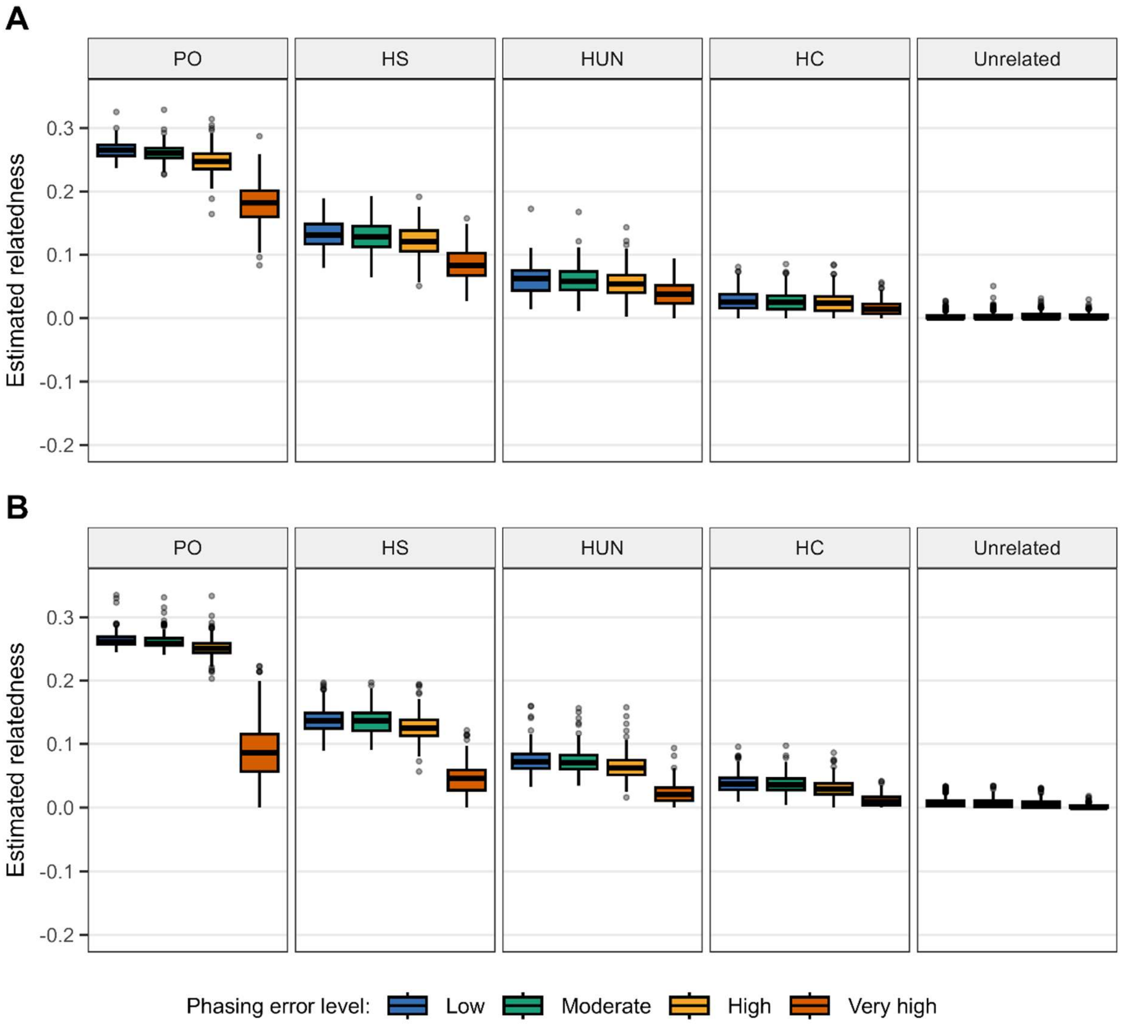

Figure S3. Estimation of recent pairwise kinship on simulated data sets and with variable levels of phasing errors. The figure shows boxplots of estimated relatedness using ZooRoH and with recent IBD only ($R_c \leq 8$), and four levels of phasing error. Low, moderate, high and very high phasing errors correspond to phasing all individuals jointly, and in subsets of 50, 20 and 10 individuals, respectively. Relatedness is reported for the following kinship categories: parent–offspring (PO), half-siblings (HS), half-uncle–nephew (HUN), half-cousins (HC) and unrelated individuals (no common ancestor in the last three generations). (**A**) Results for the first scenario, with a bottleneck. (**B**) Results for the second scenario, with a constant $N_e$.

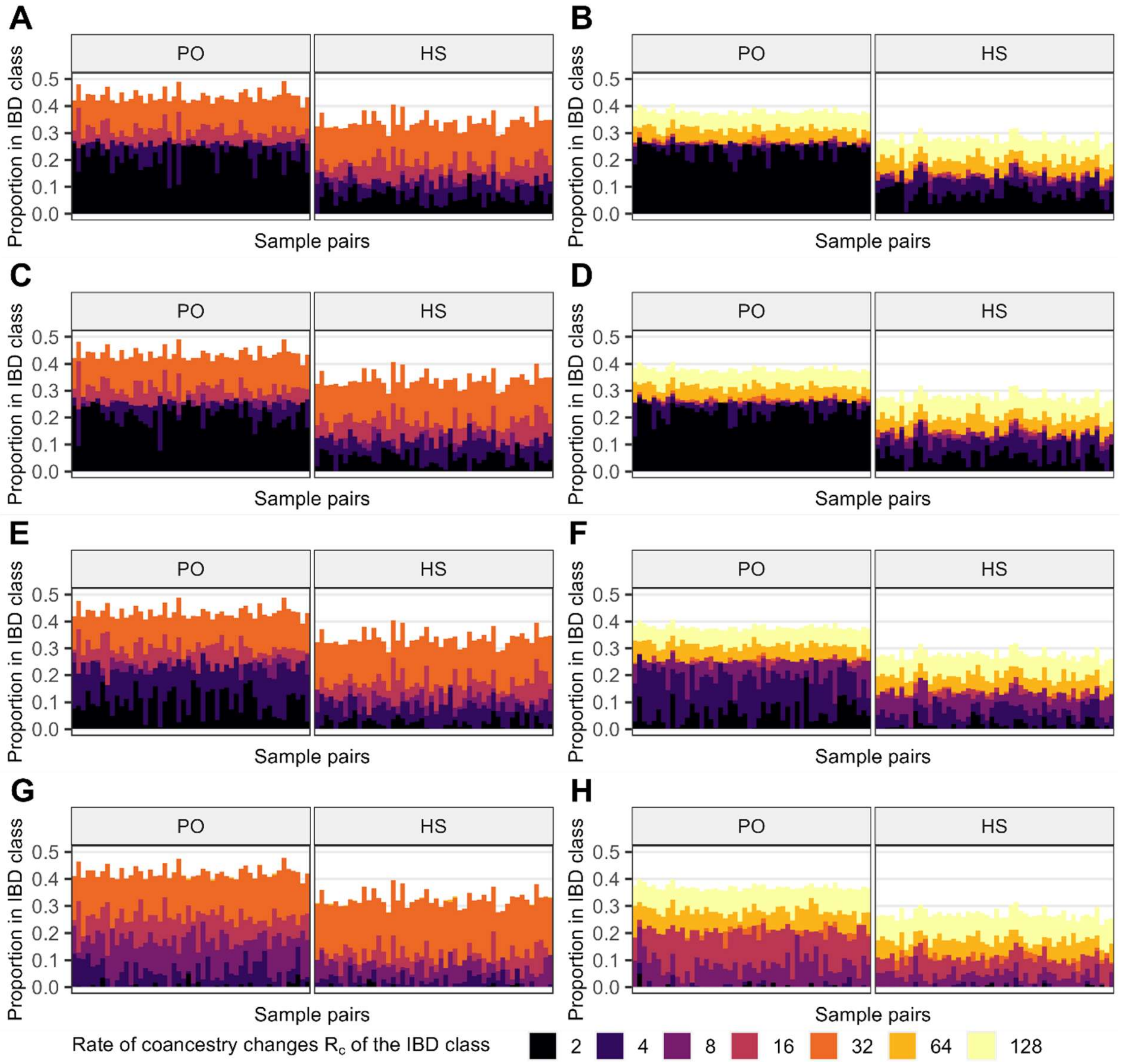

Figure S4. Estimation of pairwise kinship on simulated data sets with variable levels of phasing errors. Partitioning of the pairwise kinship into different IBD classes defined by their rates $R_C$ is reported for parent-offspring (PO) and half-siblings (HS) pairs. Results are reported for the two simulation scenarios and different levels of phasing errors. (**A**,**B**) Results for the first and second scenario, with low phasing error levels (obtained by phasing all individuals together). (**C**,**D**) Same as panels A and B, but with moderate levels of phasing errors (obtained by phasing individuals by subset of 50). (**E**,**F**) Same as panel A and B, but with high levels of phasing errors (obtained by phasing individuals by subset of 20). (**G**,**H**) Same as panel A and B, but with high levels of phasing errors (obtained by phasing individuals by subset of 10).

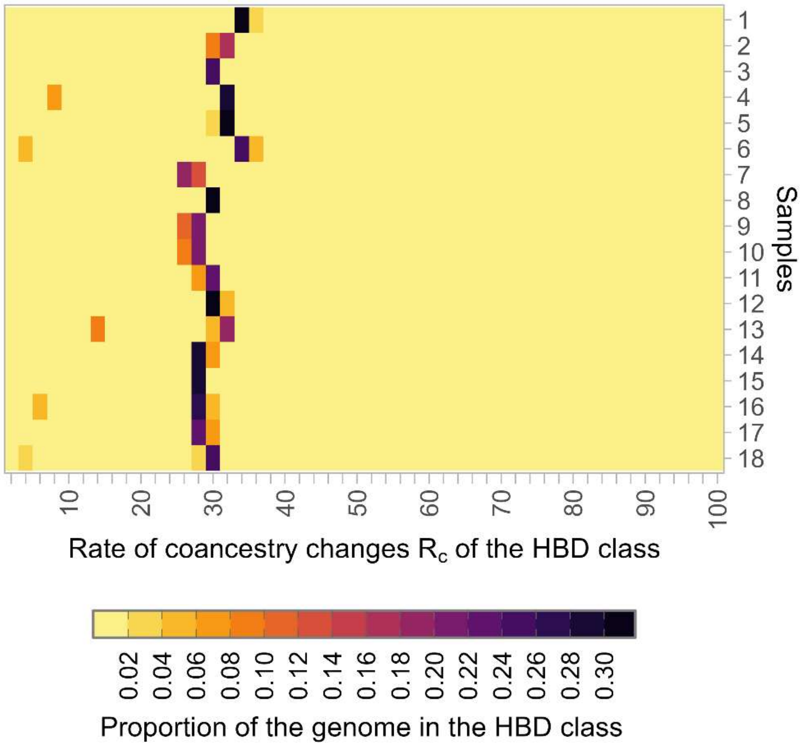

Figure S5. Individual partitioning of HBD levels with a 50-layer model in the TAF cattle population for the 18 individuals genotyped with a medium density genotyping array (23,679 polymorphic SNPs). For most individuals, only one or two HBD classes are used to capture the individual inbreeding levels.

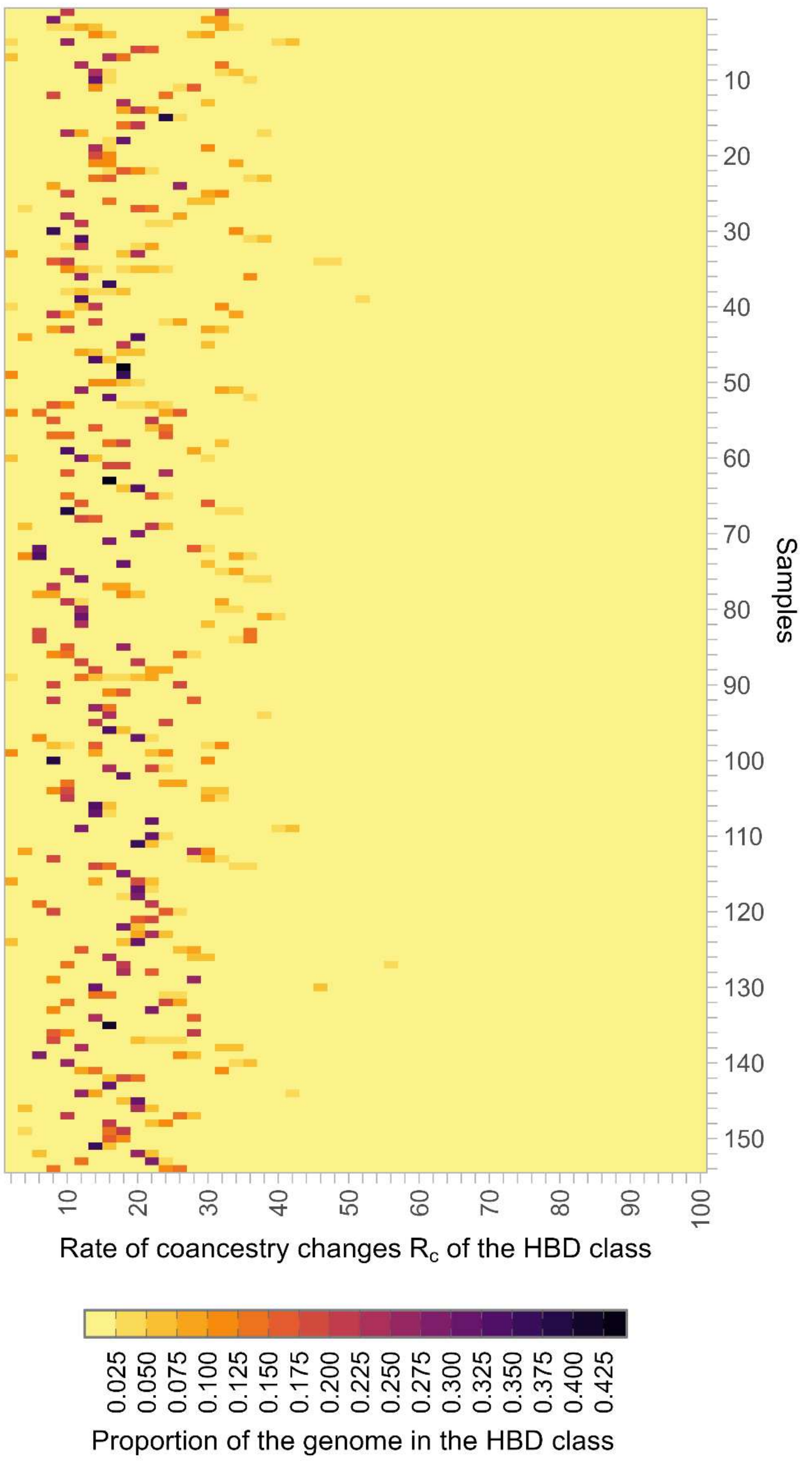

Figure S6. Individual partitioning of HBD levels with a 50-layer model in the European bison population for the 143 individuals genotyped with the BovineHD arrays (15,673 autosomal SNPs after filtering). For most individuals, only one or two HBD classes are used to capture the individual inbreeding levels.

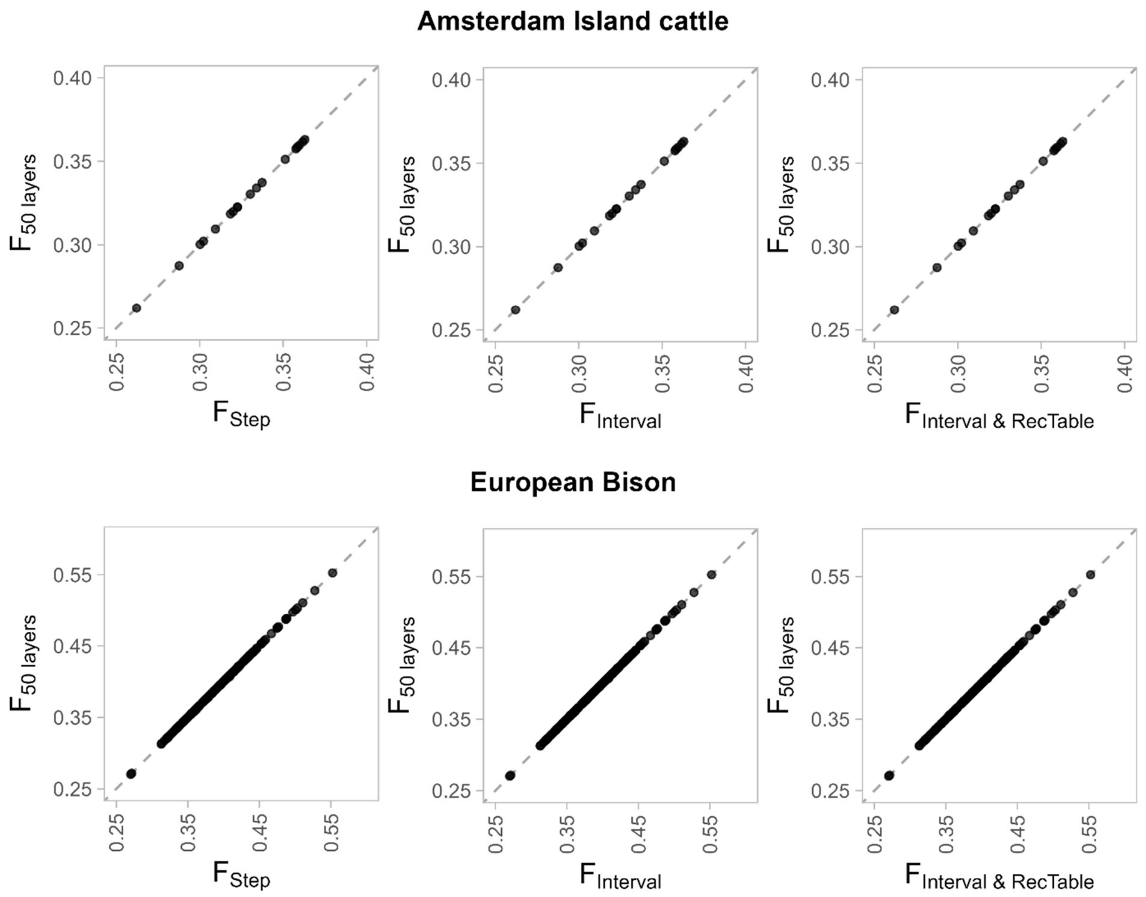

Figure S7. Comparison of inbreeding levels with different models. The inbreeding coefficients estimated using a 50-layer model were compared to those estimated using a 50-layer model and the 'Step' option ($F_{Step}$), a model based on the 'Interval' option, with ($F_{Interval\ \&\ RecTable}$) or without ($F_{Interval}$) the 'RecTable' option. The 'Step' and 'Interval' options were used to impose constant mixing coefficient for groups of five consecutive classes / generations. The model is further described in the main document. Comparisons were done for the 18 genotyped TAF individuals (Amsterdam Island cattle) and for the 143 European bison from the Lowland line.

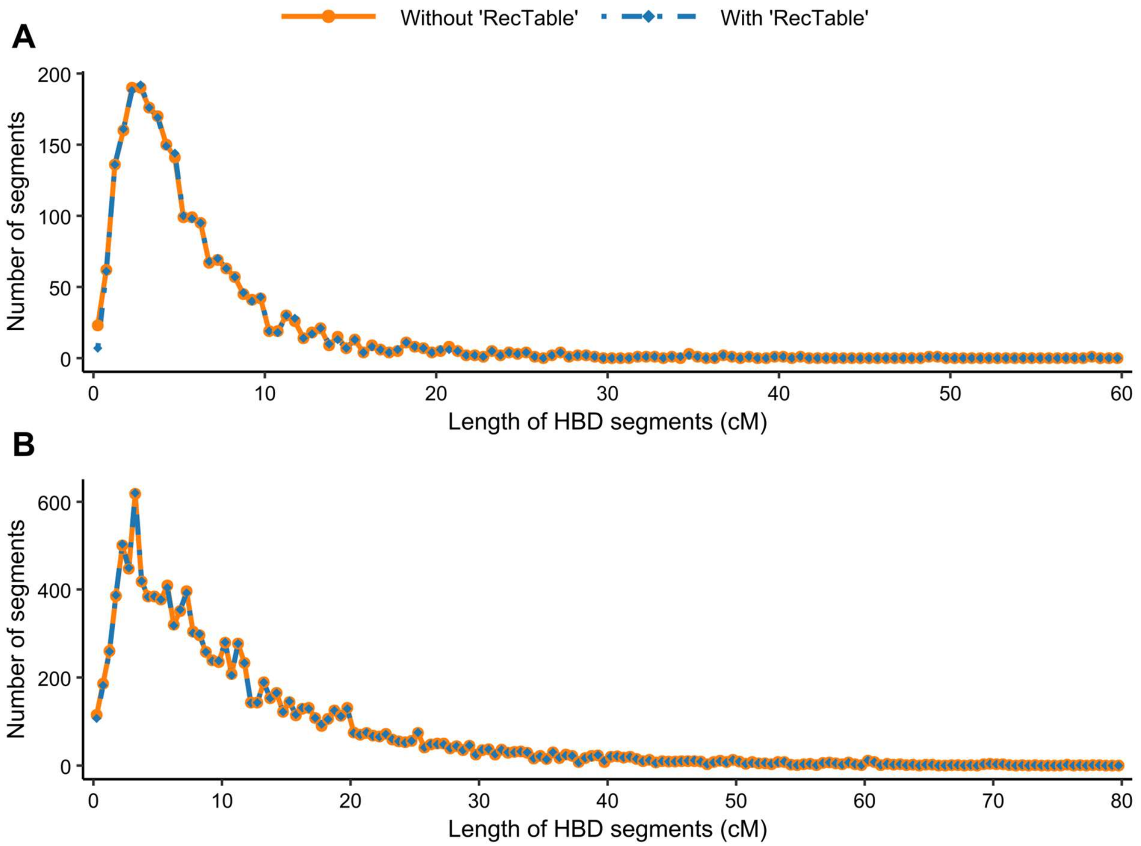

Figure S8. Distribution of HBD segment length with and without the ‘RecTable’ option. (**A**) For the 18 genotyped TAF individuals. (**B**) For the 143 European bison from the Lowland line.
